# Reprogrammed Schwann Cells Organize into Dynamic Tracks that Promote Pancreatic Cancer Invasion

**DOI:** 10.1101/2022.03.08.481473

**Authors:** Sylvie Deborde, Laxmi Gusain, Ann Powers, Andrea Marcadis, Yasong Yu, Chun- Hao Chen, Anna Frants, Elizabeth Kao, Laura Tang, Efsevia Vakiani, Annalisa Calo, Tatiana Omelchenko, Kristian R. Jessen, Boris Reva, Richard J. Wong

## Abstract

Nerves are a component of the tumor microenvironment contributing to cancer progression, but the role of cells from nerves in facilitating cancer invasion remains poorly understood. Here we show that Schwann cells (SCs) activated by cancer cells collectively function as Tumor Activated Schwann cell Tracks (TASTs) that promote cancer cell migration and invasion. Non-myelinating SCs form TASTs and have cell gene expression signatures that correlate with diminished survival in patients with pancreatic ductal adenocarcinoma. In TASTs, dynamic SCs form tracks that serve as cancer pathways and apply forces on cancer cells to enhance cancer motility. These SCs are activated by c-Jun, analogous to their reprogramming during nerve repair. This study reveals a mechanism of cancer cell invasion that co-opts a wound repair process and exploits the ability of SCs to collectively organize into tracks. These findings establish a novel paradigm of how cancer cells spread and reveal therapeutic opportunities.

**SIGNIFICANCE:** How the tumor microenvironment participates in pancreatic cancer progression is not fully understood. Here, we show that Schwann cells are activated by cancer cells and collectively organize into tracks that dynamically enable cancer invasion in a c-Jun dependent manner.

## INTRODUCTION

The migration of cancer cells and their invasion away from primary sites leads to metastasis, the principal cause of death in cancer patients. The mechanisms of cancer cell migration and invasion are complex (1, 2) and dependent on contributions from the tumor microenvironment (TME). The TME provides pro-migratory factors(2) and a matrix with distinct mechanical properties that modulate cancer cell motility(3–5). The TME cells known to affect cancer invasion through these mechanisms are immune cells and fibroblasts(4, 5). Nerves are also an important component of the TME that stimulate cancer progression. Chemical and surgical denervation experiments in animals show an inhibition of tumor growth in models of pancreatic, prostate, breast, skin, and head and neck cancers(6–11). Some cancer cells migrate, proliferate, and invade around nerves in a process called perineural invasion (PNI). Through PNI, cancer cells may spread outside the organ of the tumor’s origin, representing an unusual form of metastasis. PNI is associated with pain, paralysis, and worse patient survival(12–14). Paracrine signaling has been studied in nerve cells and a variety of secreted factors stimulate cancer cell migration(15). However, there are currently no treatments designed to inhibit PNI, cancer invasion induced by nerves, or cancer innervation(16).

Pancreatic ductal adenocarcinoma (PDAC) is an aggressive cancer with a 5-year survival rate of just 8%(17) and an incidence of PNI up to 100%(12). We demonstrated that Schwann cells (SCs), the most abundant cell type in nerves, are a mediator of PNI and facilitate pancreatic cancer cell dispersion(18). We and others have reported an increase of an SC subtype expressing elevated levels of glial fibrillary acidic protein (GFAP) when in close proximity to pancreatic cancer cells in patient specimens(18, 19). A similar increased expression of GFAP is also found in SCs responding to nerve injury. These non-myelinating repair SCs play a dynamic role in nerve regeneration and originate from GFAP-negative SCs myelinating SCs and non-myelinating (Remak) SCs expressing moderate GFAP levels that ensheathe axons in uninjured nerves(20, 21).

SCs are highly plastic cells that undergo cellular reprogramming in response to nerve injury(22–25) or infection(26). SC reprogramming during nerve repair results in the altered expression of approximately 4,000 genes, and is driven by c-Jun, Notch, Sox2, and MAPK signaling in addition to other factors (27, 28). Quiescent myelinating and Remak SCs are reprogrammed into non-myelinating nerve repair SCs that exhibit dynamic motility, release neurotrophic factors and chemokines, recruit immune cells, clear myelin by autophagy and phagocytosis, and reorganize into cellular tracks called Büngner bands. These bands are composed of elongated and aligned SCs that form a path to guide the regeneration of damaged axons(29).

Here, we found that diminished survival in patients with PDAC correlates with non-myelinating SC gene expression signatures. Cancer cells activate SCs, inducing a c-Jun dependent reprogramming that is related to that triggered by nerve injury and shifts SC differentiation state towards that of non-myelinating/repair cells. Linear tracks of such SCs, which we have named Tumor Activated Schwann cell Tracks (TASTs), function as active pathways for pancreatic cancer cell migration and invasion. This study presents a novel mechanism of cancer cell invasion that exploits dynamic cellular tracks formed by reprogrammed SCs that drive cancer migration and propagation.

## RESULTS

### Non-myelinating SC signature scores correlate with diminished survival in patients with pancreatic adenocarcinoma and with pathways related to cancer invasion

We first assessed the relationship of SCs with pancreatic cancer patient outcomes. We obtained several signatures of human SCs, including myelinating SC and non-myelinating SC signatures from the Tabula Sapiens portal using OnClass (30) (Fig. 1A). SC precursors differentiate into immature SCs that diverge to form myelinating SCs or non-myelinating (Remak) SCs. Other SCs that do not form myelin include terminal SCs and repair SCs that form from myelin and Remak cells after injury(22). We computationally derived an inferred pathway activation/suppression (IPAS) SC signature score for each available signature(31–33) (https://calina01.u.hpc.mssm.edu/pathway_assessor/) for 178 pancreatic adenocarcinoma (PAAD) patients from The Cancer Genome Atlas (TCGA) using their gene expression levels. We tested for correlations between the SC signature scores and clinical outcomes (survival and new tumor events) (Fig. 1B-E; Supplementary Fig. 1A-F). Results revealed significant correlations between clinical outcomes and non-myelinating, precursor, terminal, and global SC signature scores, with higher scores corresponding to worse outcomes (Fig. 1B-C, 1E-F; Supplementary Fig. 1A-G). There was no significant clinical correlation identified for the myelinating SC signatures (Fig. 1D,E).

**Fig. 1.**
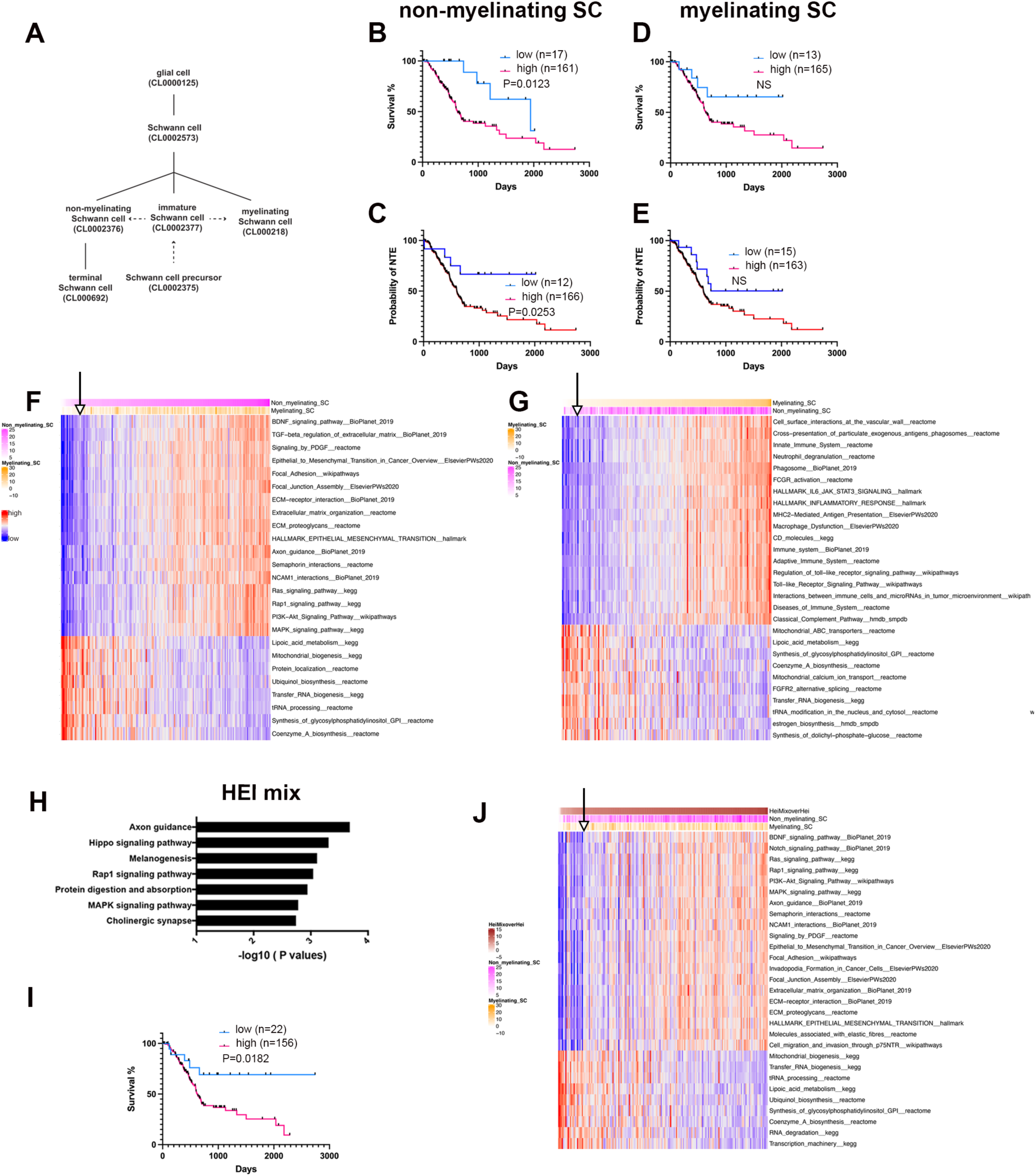
Non-myelinating SC signature scores correlate with diminished survival in patients with pancreatic adenocarcinoma and with pathways related to cancer invasion. **A**, Hierarchical organization of SCs from Tabula Sapiens (full lines) with dashed arrows indicating transitions in the SC lineage(22). **B-E**, Kaplan–Meier curves of overall survival (B,D) and new tumor events (NTE) (**C,E**) with high or low scores for signatures of non- myelinating SC (b,c), and myelinating SC (**D,E**) in 178 TCGA PAAD patients. F-G, Heatmap of gene sets correlating with high and low scores for non-myelinating SC signature (F) and myelinating SC signature (G) in TCGA PAAD patients. Columns represent TCGA PAAD samples that have been rank ordered by the top row signature. Arrows indicate survival cutoff used in b and d with low survival patients at the left of the arrow. **H**, Top 7 enriched pathways in HEI-286 co-cultured with MiaPaCa-2 as compared with HEI-286 SCs alone (EnrichR, human KEGG 2019 dataset). **I**, Kaplan–Meier curve of overall survival with high or low scores for the cancer exposed HEI-286 SC (HEImix) signature in 178 TCGA PAAD patients. **J**, Heatmap of gene sets correlating with high and low scores for cancer exposed HEI-286 SC (HEImix) signature in TCGA PAAD patients. Columns represent TCGA PAAD samples that have been rank ordered by the top row signature.The arrow indicate survival cutoff used in i with low survival patients at the left of the arrow.

Further analysis indicated that the gene expression signature of non-myelinating SCs positively correlated with multiple pathways related to cancer invasion in the dataset from TCGA, including epithelial–mesenchymal transition, MAPK signaling, PI3-Akt signaling, extracellular matrix organization, and others. The non-myelinating SC signature also correlated with axonal guidance gene sets (Fig. 1F). The myelinating SC signature positively correlated with genes sets related to immune cells and immune function (Fig. 1F; Supplementary Fig. 1H,I). The negatively correlated gene sets were similar for both non-myelinating and myelinating signatures and involved pathways with lipoic acid metabolism, mitochondrial activity, and tRNA processing (Fig. 1F,G).

We created a cancer cell–exposed SC signature (HEI-mix) experimentally using the non-neoplastic human SC line HEI-286 that was co-cultured either with or without MiaPaCa- 2 pancreatic cancer cells. GO pathway analysis of the transcriptome of SCs sorted by FACS after cancer cell co-culture revealed a significant enrichment of genes related to axon guidance and MAPK signaling pathways (Fig. 1H). We then tested correlations between our cancer cell–exposed SC signature score (HEI-mix) with clinical outcomes using the dataset from TCGA. HEI-mix scores were significantly correlated with patient survival, with higher scores corresponding to worse patient survival (Fig. 1I). Furthermore, the gene sets and pathways that correlated the most with this signature were nearly identical to those that correlated with the non-myelinating SC signature and are linked to cancer invasion (Fig. 1J). Among all SC signatures tested, the non-myelinating SC signature was the cell signature that most closely correlated with the HEI mix signature (Supplementary Fig. 1J). Altogether, these findings link both non-myelinating SCs and SCs whose transcriptome profile has been altered by the presence of cancer cells with a worse clinical severity of PAAD and transcriptomic profiles relating to cancer cell invasion and axon guidance.

### SCs wrap and align cancer cells *in vivo* and *in vitro*

We have previously reported the presence of non-myelinating GFAP^+^ SCs in PDAC (18). To explore the role of these cells in cancer invasion, we examined their distribution in human specimens of PDAC. GFAP^+^ SCs closely associated with cancer cell clusters in the nerves with PNI (Fig. 2A-H). Nerves invaded by cancer cells exhibit SC heterogenicity in GFAP expression, with strongly GFAP^+^ SCs located near the cancer cells (Fig. 2B-D; Supplementary Fig. 2A). The GFAP^+^ SCs appeared to align the cancer cells within the nerves (Fig. 2E-H). Orthogonal views (xy and xz) of confocal images revealed that GFAP^+^ SCs enveloped the cancer cells (Fig. 2H). In PDAC stroma rich with nerve fibers, we also found that GFAP^+^ SCs formed close associations with cancer cells. Sometimes a GFAP^+^ SC encircled a single cancer cell, or a small cluster of cancer cells (Fig. 2I-K). Other times, columns of cancer cells were found intertwined with SCs insinuating around them (4 out of 6 patients with PNI) (Fig. 2L,M; Supplementary Fig. 2B). We examined SC and cancer cell interactions using a PNI model in which Panc02 cancer cells injected into sciatic nerves of mice migrate toward the spinal cord(34, 35). As in PNI human samples, longitudinal sections of the mouse sciatic nerves revealed that GFAP^+^ SCs surrounded the clusters of cancer cells (Fig. 2N,O).

**Fig. 2.**
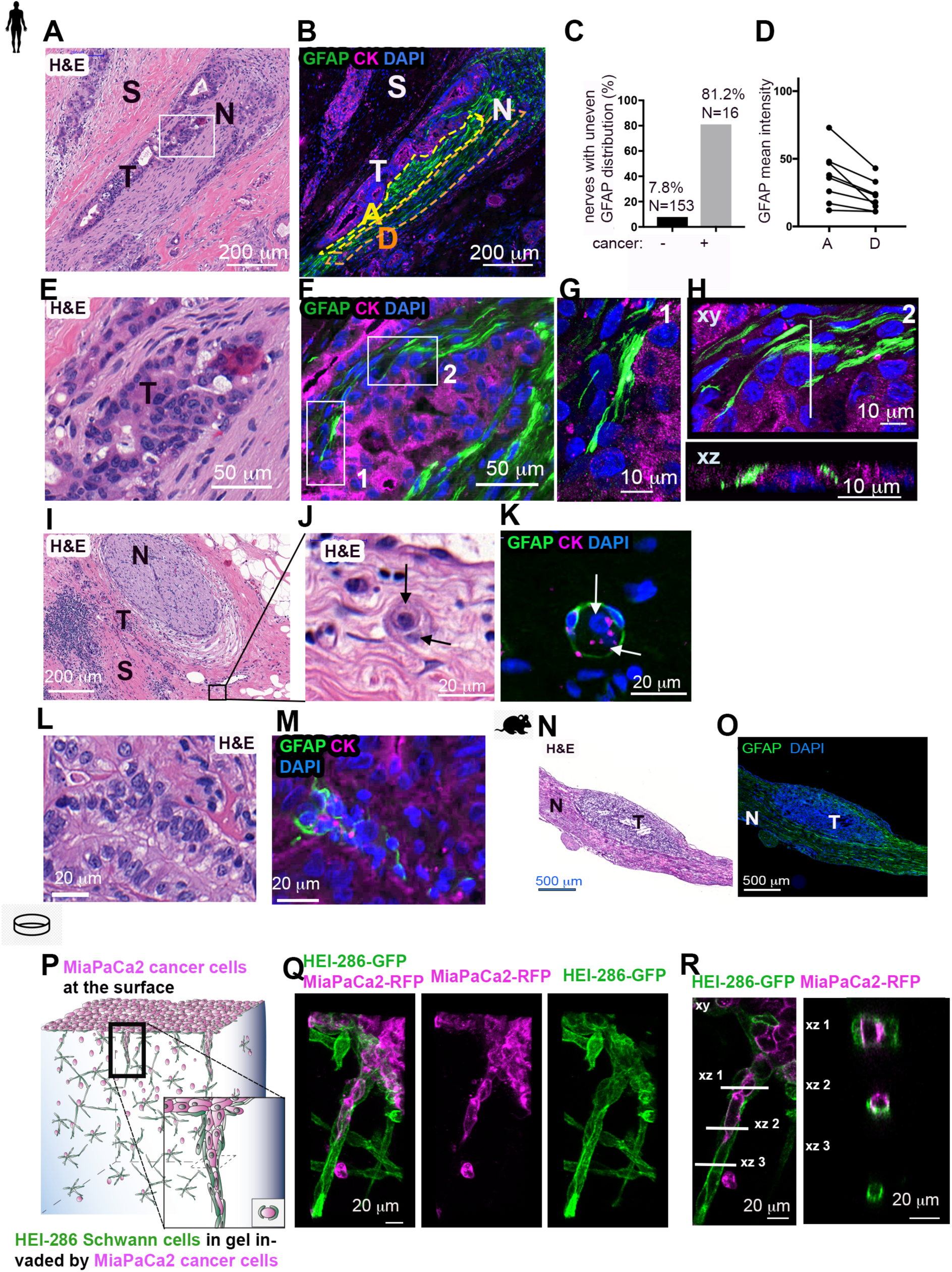
SCs wrap and align cancer cells. **A**, H&E section of a human PDAC specimen with PNI. Scale bar, 200 μm. T, tumor; N, nerve; S, stroma. **B**, GFAP (green) and cytokeratin (CK, magenta) staining in a section adjacent to (**A**) showing cancer cells surrounded by GFAP^+^ SCs in a nerve. The yellow and orange dotted regions of the nerve are adjacent (A) or distal (D) to the cancer cells. Scale bar, 200 μm. **C**, Quantification of nerves with an uneven GFAP distribution comparing when cancer is visibly present or absent. **D**, Quantification of GFAP mean intensity in adjacent (A) and distal (D) regions of nerves. **E**, Enlargement of rectangle in (**A**). Scale bar, 50 μm. **F**, GFAP and CK staining of an adjacent section of (**E**). Scale bar, 50 μm. **G**, Confocal images corresponding to rectangle 1 in (**f**). Scale bar, 10 μm. **H**, Confocal images corresponding to rectangle 2 in (**F**). Top image is an xy maximum projection and bottom image is an xz image corresponding to the dotted line. Scale bars, 10 μm. **I**, H&E section of a human PDAC specimen with a nerve (N) and tumor cells (T) in the neighboring stroma (S). Scale bar, 200 μm. **J**, Enlargement of (**i**). Arrows indicate 2 cancer cells. Scale bar, 20 μm. **K**, GFAP and CK staining of a section adjacent to (**J**) showing GFAP^+^ SCs wrapping two cancer cells (arrows). Scale bar, 20 μm. **L-M**, H&E staining and adjacent GFAP and CK staining of a human PDAC specimen showing GFAP+ SCs around aligned cancer cells. Scale bars, 20 μm. **N-O**, H&E staining and adjacent GFAP staining of a longitudinal section of a murine sciatic nerve injected with cancer cells. Scale bars, 500 μm. **P**, Schematic representation of HEI-286 SCs and MiaPaCa-2 cancer cells in a 3D Matrigel invasion assay. The MiaPaCa-2-RFP cancer cells are placed on the surface of a Matrigel chamber in which HEI-286 SCs have grown. Cancer cells invade into the gel as a chain of cancer cells surrounded by HEI-286 SC tracks. **Q**, Maximum projection of confocal images showing chain of cancer cells lined up within a tubular structure of HEI-286 SCs in Matrigel. Scale bar, 20 μm. **r**, Single focal planes of confocal images showing MiaPaCa-2 cancer cells and HEI-286 SCs organized into columns. Longitudinal (left) and transverse (right) images, correspond to the indicated positions. Scale bars, 20 μm.

We next examined SC and cancer cell interactions with an *in vitro* 3D model of cancer invasion to assess if SCs wrap cancer cells. HEI-286 SCs expressing GFP facilitate the downward invasion of a red fluorescent human pancreatic cancer cells MiaPaCa-2 seeded on top of Matrigel chambers(18). Three days after the addition of MiaPaCa-2, HEI-286 SCs were found closely associated with chains of cancer cells (Fig. 2P,R). HEI-286 SCs wrapped around and aligned with chains of MiaPaCa-2 cancer cells (Fig. 2Q,R), creating invasive columns of cells. In contrast, MiaPaCa-2 cells seeded on Matrigel chambers lacking HEI-286 SCs, or in the presence of NIH3T3 fibroblasts, failed to invade and did not organize into chains (Supplementary Fig. 2C,D). Collectively, the *in vivo* and *in vitro* images revealed a consistent pattern of organization in which SCs wrapped cancer cells and created columns of SCs and cancer cells. These observations suggest that SCs reorganize cancer cells into chains and create tracks for migration, which we have named Tumor Activated Schwann cell Tracks (TASTs).

### SCs organize into dynamic tracks that allow cancer cell migration

We used microfabricated channels to investigate TASTs. They allow the alignment of HEI- 286 SCs in 3D as in TASTs and the visualization of cell motility and cell behavior. To determine if cancer cells migrate along TASTs, we assessed dynamic interactions between HEI-286 SCs and MiaPaCa-2. HEI-286 SCs and MiaPaCa-2 cells, seeded together in wells, migrate into the channels and cells organized into columns of SCs and cancer cells (Fig. 3A,B). Confocal microscopy showed that HEI-286 and MiaPaCa2 cells occupied the full width of the 10–20 μm wide channels (Supplementary Fig. 3A, xz 2 and xz 4). HEI-286 SCs wrapped around MiaPaCa2 cells at 80% of contact sites, either partially (Supplementary Fig. 3A, xz 3) or completely (Fig. 3B, xz). This wrapping behavior around centrally positioned cancer cells mirrored observations from our 3D Matrigel invasion assay (Fig. 2P,R) and PDAC sample (Fig. 2H,K). We examined cancer cell dynamics in the 3D microchannels seeded first with HEI-286 SCs (Supplementary Fig. 3B). Cancer cells migrated within the microchannels lined by HEI-286 SCs (Fig. 3C; Supplementary Video 1). HEI-286 wrapped themselves around MiaPaCa-2, lined themselves along the microchannel walls, and permitted cancer cell migration through the microchannel (Fig. 3C,D; Supplementary Video 2).

**Fig. 3.**
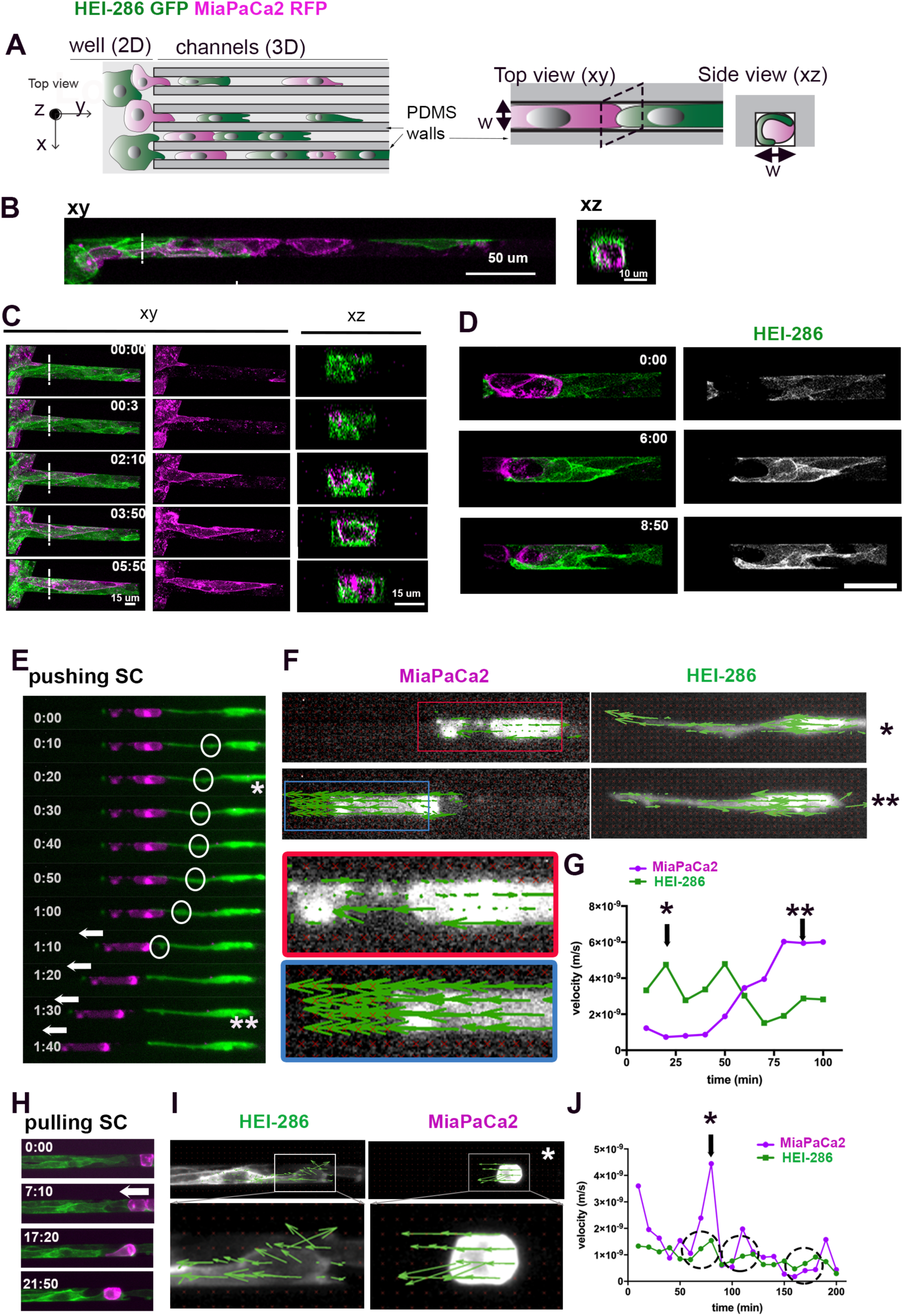
SCs form dynamic tracks for cancer cells. **A**, Schematic of microchannels with HEI-286 SCs (green) and MiaPaCa2 (magenta). Both cell types are seeded in the adjacent well (left, 2D) and enter microchannels (3D) where they make contact with each other. **B**, Confocal images of HEI-286 SCs and MiaPaCa2 within microchannels in longitudinal (xy) and transverse sections (xz). Scale bars, 50 μm. **C**, Confocal images of time-lapse movies showing a cancer cell moving in a microchannel lined by HEI-286 SCs. Time is h:min. Scale bars, 15 μm. **D**, Confocal images of time-lapse movies showing a HEI-286 SC wrapping around a cancer cell. Scale bar, 50 μm. **E**, Fluorescent images of time-lapse movie showing a HEI-286 SC pushing a cancer cell. Arrows indicate cancer cell displacement. Circles indicate intracellular movement within the SC (* and ** are time points shown in E). **F**, Fluorescent images of a HEI-286 SC and cancer cell from (e) at two time points (* and **) overlaid with vectors obtained by PIV analysis indicating vector direction. **G**, Quantification of the mean instantaneous velocity of the HEI-286 SC and cancer cell in (**E**), (**F**), and (**G**). * and ** indicate corresponding time points. **H**, Fluorescent images of time-lapse movie showing HEI- 286-SCs pulling a cancer cell. Arrows indicate cancer cell displacement. **I**, Fluorescent images of the HEI-286 SCs and cancer cell from (**H**) at one time point (*) overlaid with vectors obtained by PIV analysis. **J**, Quantification of the mean instantaneous velocity. (*) indicates corresponding time point in (**I**). Dotted circles indicate periods with synchronized increases in velocity for both cancer cells and SCs.

### SCs exert forces on cancer cells that propel migration

To assess if SCs influence the movement of cancer cells within TASTs, we analyzed time-lapse microscopy and particle image velocimetry (PIV). A single HEI-286 SC was able to propel a cancer cell through a pushing effect (Fig. 3E; Supplementary Video 3). Circles show a propulsive wave within a HEI-286 SC that propagated from the center of the cell toward its contact point with a static cancer cell. The wave then created a SC protrusion at the site of contact, which propelled the cancer cell away from the HEI-286 SC. PIV analysis of both cells revealed intracellular movement corresponding to an intracellular propulsive wave within the SC prior to the cancer cell displacement (Fig. 3F,G), with movement of both cells in the same direction (Fig. 3G). We also observed squeezing motions by HEI-286 SCs surrounding a MiaPaCa-2 cell. The combined effect of wrapping and squeezing of the MiaPaCa-2 cells led to propulsive forces moving the cancer cell (Supplementary Fig. 3C; Supplementary Video 4). HEI-286 SCs also exerted pulling forces, making contact with MiaPaCa-2 cells and then retracting the cancer cells toward themselves (Fig. 3H; Supplementary Video 5). PIV analysis revealed that HEI-286 SCs and cancer cells displayed coordinated directional vectors (Fig. 3I) and instantaneous velocity intensities (Fig. 3J).

Increases in SC velocities correlated temporally with increases in cancer cell velocities (circles in Fig. 3J). The HEI-286 SCs that formed the TASTs permitted MiaPaCa-2 cells to pass through them centrally and directly applied forces on the cancer cells. HEI-286 SCs were able to push, squeeze, or pull cancer cells, contributing to cancer cell motility.

### Cancer cells activate c-Jun in SCs

Following nerve trauma, myelinating and Remak SCs trans-differentiate into SCs that actively engage in nerve repair. This transition is controlled in part by the transcription factor c-Jun(27), which reprograms myelinating and Remak SCs into SCs that align into Büngner bands to enable axonal guidance and nerve regeneration. This SC organization during nerve repair shows similarities to our observations of SC alignment into tracks enabling cancer cell migration *in vitro*. In the PAAD TCGA dataset, we identified a significant correlation between JUN expression level and the non-myelinating SC signature (p=0.0004 r=0.2608), but not with the myelinating signature (p=0.8334, r=0.01588). Furthermore, higher JUN expression correlated with worse patient survival (Fig. 4A, p=0.035).

**Fig. 4.**
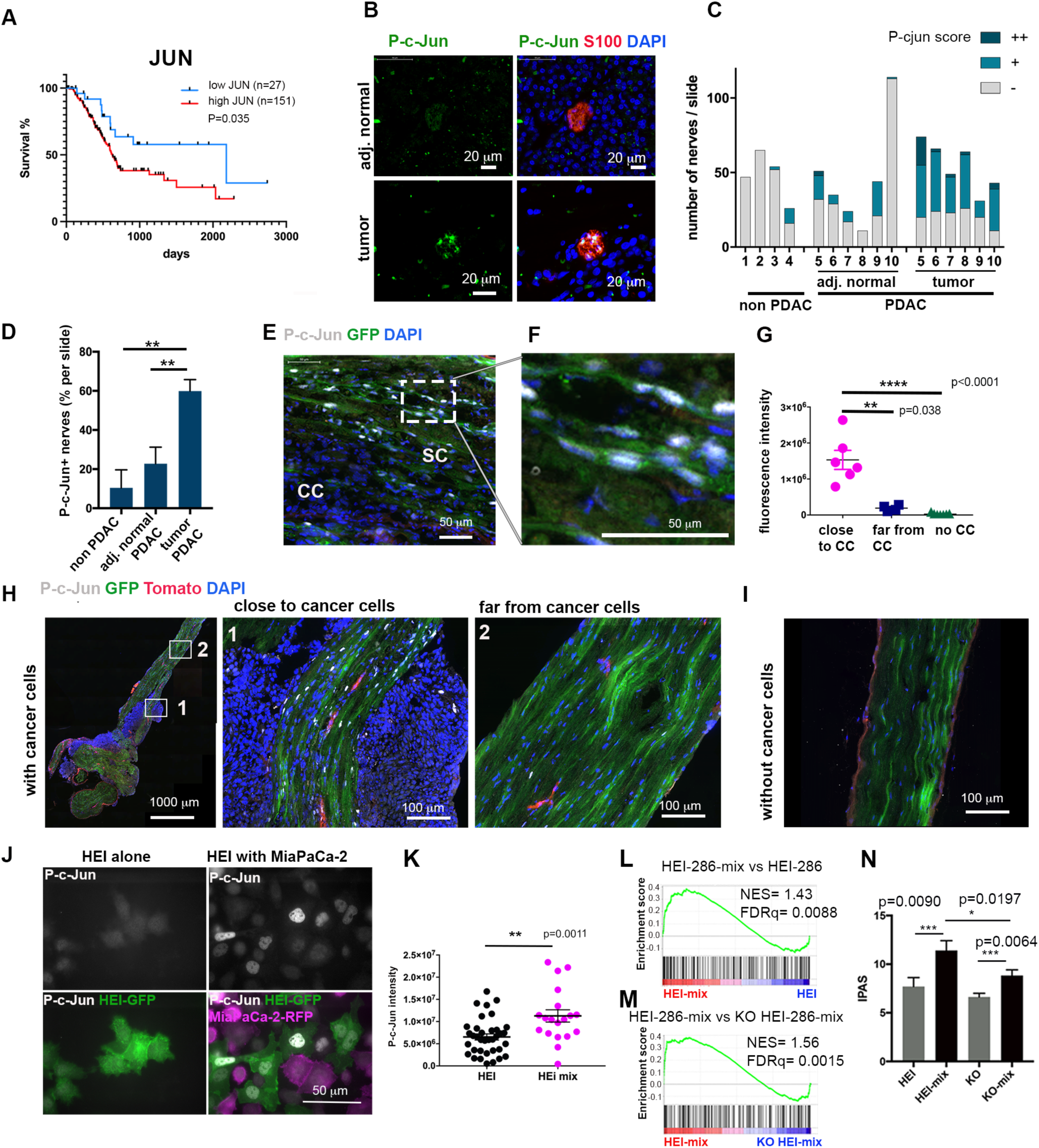
Cancer cells induce SC c-Jun activation and reprogramming. **A**, Kaplan–Meier curve of overall survival (OS) with high or low JUN expression in TCGA PAAD patients.**B**, P-c-Jun staining in S100-labeled nerves from PDAC specimens that are close to tumor as compared with nerves from adjacent normal tissue. Scale bars, 20 μm. **C,D**, Assessment of nerves from within pancreatic specimens of non-PDAC pathology, PDAC, and the normal tissues adjacent to PDAC. **C**, Quantification of number of nerves per slide with no (-), low (+), or high (++) intensity P-c-Jun staining. Each x axis number represents a patient. **D**, Percentage of P-c-Jun positive nerves (scored + or ++) in the pancreatic specimens of non- PDAC pathology, PDAC, and the normal tissues adjacent to PDAC (non-PDAC, n=4; PDAC and adjacent normal, n=6 each; mean ± SEM). **E,F**, Immunofluorescence of P-c-Jun (white) in a Panc02 injected murine sciatic nerve expressing GFP^+^ SCs. P-c-Jun is expressed in the green SCs but not in the Panc02 cancer cells (CC). Scale bars, 50 μm. **G**, Quantification of P- c-Jun fluorescence intensity in SCs of uninjected nerves, and in SCs close to or far from injected Panc-02 tumor cells (near tumor: n=6; far from tumor: n=4, PBS-injected: n=7, mean ± SEM, representative of three independent experiments). Fluorescent intensity is the mean intensity measured per area of nerve covered by green SCs. **H**, P-c-Jun staining in a sciatic nerve injected with Panc-02. (1) Region adjacent to the tumor. (2) Region far from the tumor. Scale bars, 1000 and 100 μm. **I**, P-c-Jun staining in a normal nerve without cancer cells. Scale bar, 100 μm. **J**, Immunofluorescence of P-c-Jun in HEI-286 GFP SCs grown alone or grown mixed with MiaPaCa-2 RFP. Top images show P-c-Jun staining alone in HEI-286 SCs alone (left) and co-cultured HEI-286 mixed with MiaPaCa-2 (right). Bottom overlay images allow identification of HEI-286 SCs (green) and MiaPaCa2 (magenta). Scale bar, 50 μm. **K**, Quantification of P-c-Jun fluorescence intensity in HEI-286 SCs grown alone or mixed with MiaPaCa2. Mean fluorescence intensity was measured per HEI-286 SC (n>15 cells/group, mean ± SEM, representative of three independent experiments). **L-M**, Gene set enrichment analysis (GSEA) assessing axon guidance genes in cancer co-cultured HEI-286 SCs compared with HEI-286 SCs alone (**L**) and in cancer co-cultured HEI-286 compared with cancer co- cultured c-Jun KO HEI-286 SCs (**M**). NES is normalized enrichment score. **N**, Inferred Pathway Activation/Suppression (IPAS) scores for the axon guidance genes (KEGG 2019).

To explore whether c-Jun-dependent reprogramming occurs during SC and cancer cell interactions, we assessed c-Jun phosphorylation in the nuclei of SCs in proximity to cancer cells in human surgical specimens. We identified strong P-c-Jun expression in nerves within PDAC, reduced P-c-Jun expression in nerves in normal tissue adjacent to PDAC, and minimal P-c-Jun expression in nerves from non-PDAC pancreatic specimens (Fig. 4B-D). This pattern suggests that cancer cells trigger SC c-Jun activity.

We also assessed SC P-c-Jun expression in the murine model of nerve invasion (Fig. 2 N,O) using transgenic P0-CRE mT/mG mice with GFP expressing SC after injecting Panc02 cancer cells into the sciatic nerves. P-c-Jun localized in GFP expressing SCs but not in cancer cells (Fig. 4D,E). We assessed P-c-Jun expression in SCs that were: (1) close to the cancer cells, (2) far from the cancer cells, or (3) in non-cancer bearing mice. Immunofluorescence staining revealed a significant increase of SC P-c-Jun staining only when in close proximity to the cancer cells (Fig. 4F-H).

We tested if SC c-Jun activation could be mediated directly by cancer cells in the absence of the TME using HEI-286 SCs expressing GFP co-cultured with MiaPaCa-2 cells expressing RFP. P-c-Jun expression was significantly increased in the HEI-286 cells mixed with MiaPaCa-2-RFP as compared with HEI-286 cells alone (Fig. 4I,J). These results demonstrate that cancer cells directly induce SC c-Jun activation.

To assess the transcriptional effects of cancer cell–mediated SC c-Jun activation, we generated c-Jun knockout (KO) HEI-286 SCs (Supplementary Fig. 4A-C). We examined the gene expression profiles of c-Jun KO HEI-286 SCs co-cultured with MiaPaCa-2 cancer cells as compared with c-Jun KO HEI-286-SCs grown alone, and derived the cancer-exposed c-Jun KO SC signature (KO-HEI-mix). The correlation of this signature score with patient survival was less significant (p=0.0480) than the correlation of the control HEI-mix signature score with patient survival (p=0.0182, Fig. 1I). In HEI-286 cells, GO pathway analysis and gene set enrichment analysis (GSEA) revealed enrichment of axon guidance genes in co-cultured HEI- 286 SCs as compared with the HEI-286 SCs grown alone (Fig. 1H and Fig. 4K) and other c-Jun-related pathways such as the MAPK signaling pathway. To assess the relevance of c-Jun in SC axon guidance activation by cancer, we examined the gene expression profile of HEI- 286 SCs as compared with c-Jun KO HEI-286 SCs co-cultured with MiaPaCa-2 cells. GSEA revealed an enrichment in axon guidance genes for HEI-286 SCs with intact c-Jun (NES=1.56, FDRq =0.0015) (Fig. 4L). The IPAS scores of axon guidance genes (KEGG- 2019) for the different conditions revealed the comparative enrichment to be HEI_mix>KO_mix>HEI>KO (Fig. 4M). Both of these analyses implicate a role for SC c-Jun in the regulation of axon guidance gene expression following cancer exposure.

SC gene expression changes during nerve repair that are dependent on c-Jun have been identified(27). We tested enrichment of the gene set upregulated in injured sciatic nerves of wild type (WT) as compared with c-Jun-KO mice(27) in our co-cultured SCs. GSEA revealed an enrichment of c-Jun nerve repair genes in co-cultured HEI-286 SCs (FDRq<0.25) as compared with HEI-286 SCs grown alone (NES=1.191, FDRq =0.188) (Supplementary Fig. 4E) despite species differences. Similarly, IPAS revealed a higher enrichment of c-Jun nerve repair genes in the co-cultured HEI-286 SCs as compared with HEI-286 SCs grown alone (Supplementary Fig. 4F). We then tested whether our identified genes upregulated in co-cultured HEI-286 SCs as compared with co-cultured c-Jun KO HEI-286 SCs (452 genes with FC>2, p<0.05) were enriched in the injured (cut) or intact (uncut) murine sciatic nerve from the same study(27). GSEA revealed an enrichment of these human genes in the murine- injured sciatic nerves (NES=1.318, FDRq=0.015) (Supplementary Fig. 4G). These findings indicate that transcriptomic changes in HEI-286 SCs following cancer exposure show parallels with SC reprogramming following nerve injury. Both axon guidance and c-Jun-related nerve repair gene sets in SCs are induced by cancer.

### SC c-Jun enhances cancer migration along TASTs in microchannels

We assessed how SC c-Jun impacts cancer cell migration within microchannels. The c-Jun KO HEI-286 SCs did not migrate as fast as control HEI-286 SCs (Supplementary Fig. 5A-C), made shorter columns than control HEI-286 SCs when combined with MiaPaCa-2 (Supplementary Fig. 5D), and were less able to wrap cancer cells. While 80% of control HEI- 286 SCs wrapped cancer cells at site of contact as described (Fig. 3A,B), only about 50% of the c-Jun KO HEI-286 SCs wrapped cancer cells (Fig. 5A,B). The c-Jun KO HEI-286 SCs sometimes clustered in the central part of the microchannel, surrounded by cancer cells that lined the periphery (Fig. 5A).

**Fig. 5.**
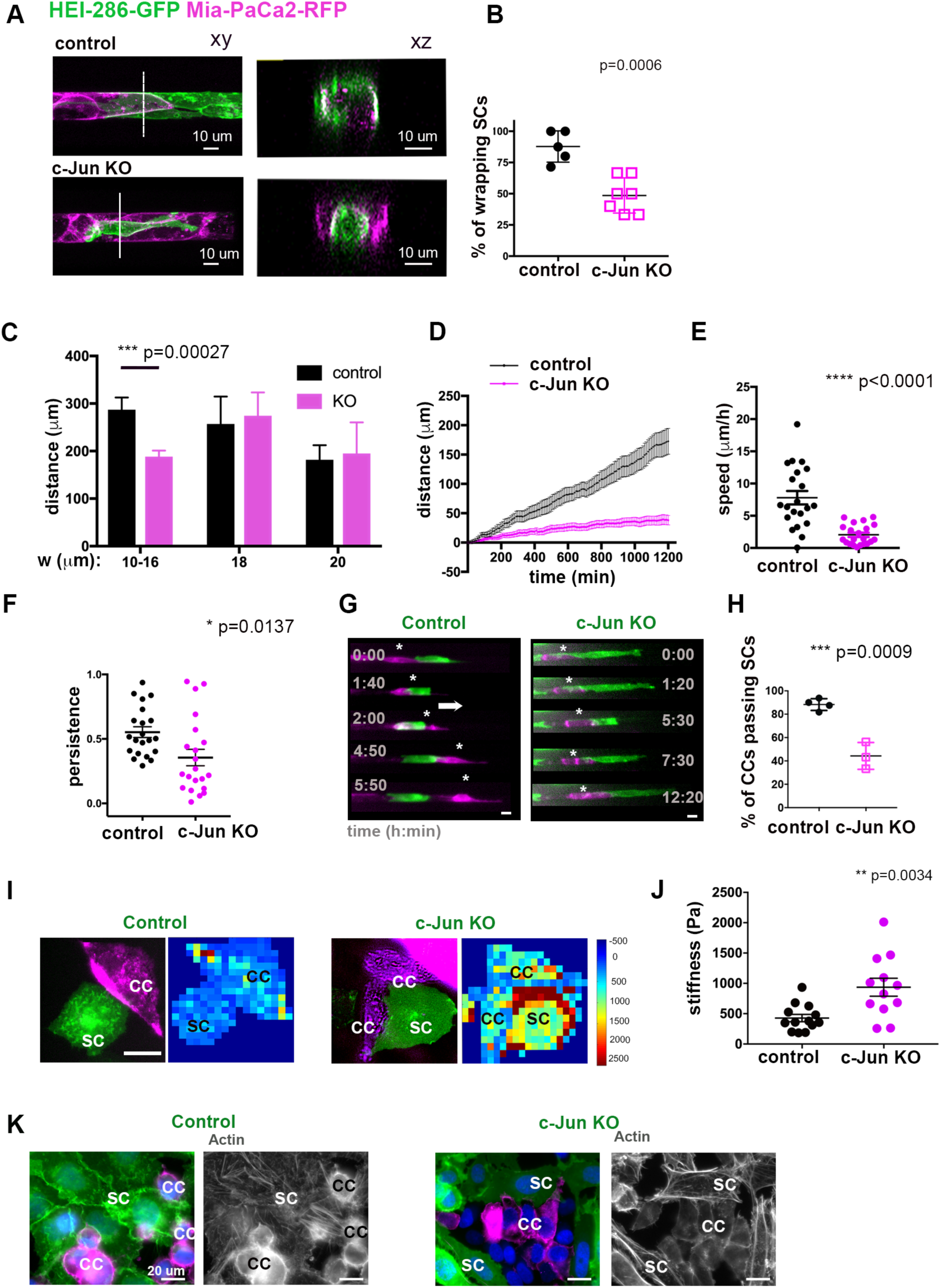
SC c-Jun faciliates cancer migration along SC tracks in microchannels. **A**, Confocal images of HEI-286 with MiaPaCa2 (top), and c-Jun-KO HEI-286 with MiaPaCa2 (bottom) within microchannels showing longitudinal (xy) and transverse sections (xz). Scale bars, 10 μm. **B**, Quantification of percentage of HEI-286 SCs wrapping around MiaPaCa-2 cells for control or c-Jun KO HEI-286 SCs (n=5-7 recordings per group with a total of 23-27 cells/group, mean ± SEM). **C**, Quantification of distance migrated by cancer cells in microchannels of different widths, and occupied by HEI-286 SCs (n=9-20 cells per channel size). **D**, Tracks of cancer cells in microchannels occupied by control and c-Jun KO HEI-286- SCs Average representation of 21 MiaPaCa-2 cells in each group (F test with F=930.1 DFn=1 DFd=4980 p<0.0001). **E**, Quantification of cancer cell speed (absolute values) in microchannels occupied by control vs. c-Jun KO HEI-286 SCs (n=21 cells in each group). **F**, Quantification of MiaPaCa-2 cell directional persistence (absolute values) in microchannels occupied by control vs. c-Jun KO SCs HEI-286 SCs (n=21 cells in each group). **G**, Fluorescent images of time-lapse movies showing the behavior of a MiaPaCa-2 after HEI-286 SC contact. A cancer cell passes by a control HEI-286 SC, while another is blocked by a c- Jun KO HEI-286 SC. Scale bars, 15 μm. **H**, Quantification of (G) showing the percentage of cancer cells passing by a HEI-286 SC (n=3 experiments per group with at least 11 cells/group in each experiment, mean ± SEM). **I**, Images showing stiffness maps of co-cultured HEI-286 SCs vs. c-Jun KO HEI-286 SCs. Scale bar, 15 μm. **J**, Quantification of stiffness of HEI-286 SCs vs. c-Jun KO HEI-286 SCs, measured by atomic force microscopy (n=12-13 cells/group, mean ± SEM, representative of two independent experiments). **K**, Images showing actin staining in co-cultured HEI-286 SCs vs. c-Jun KO HEI-286 SCs. Scale bar, 20 μm.

When HEI-286 SCs were seeded first in 10–16 µm-wide microchannels, MiaPaCa-2 migrated significantly longer distances combined with control HEI-286 SCs as compared with c-Jun KO HEI-286 SCs (Fig. 5C). Interestingly, this difference was not seen in channels of wider diameter (Fig. 5C), showing that HEI-286-SC c-Jun-dependent enhancement of MiaPaCa-2 migration occurs only under confinement conditions.

We tracked individual MiaPaCa-2 cell movement within microchannels of 10-16 µm width occupied by HEI-286 SCs to assess migration dynamics over 20 hours. Cancer cells displayed persistent migration and reached an average speed of 7.80±1.04 µm/h in the presence of control HEI-286 SCs. In contrast, the average cancer cell speed was significantly reduced at 2.04±0.34 µm/h in presence of c-Jun KO HEI-286 SCs (Fig. 5D,E; Supplementary Fig. 5E). The average directional persistence of cancer cell migration was significantly greater in the microchannels seeded with control HEI-286 SCs as compared with c-Jun KO HEI-286 SCs (Fig. 5F). MiaPaCa-2 often either failed to move, or moved backwards, in presence of the c-Jun KO HEI-286 SCs (Supplementary Fig. 5E).

We next evaluated the ability of a migrating cancer cell to either pass or be blocked by HEI-286 SCs in a microchannel after making contact. Our experiments demonstrated that control HEI-286 SCs enabled the passage of cancer cells in a confined microchannel (about 80% of the time after contact), while c-Jun KO HEI-286 SCs were prone to blocking cancer cell passage (about 40% of the time) (Fig. 5G,H; Supplementary Video 6).

To test whether differences in the physical properties of control as compared with c-Jun KO HEI-286 SCs contribute to cancer cell transit, we measured the physical stiffness of HEI-286 SCs using atomic force microscopy (AFM). We reasoned that softer and more pliable HEI-286 SCs might facilitate the passage of MiaPaCa-2 cells, by being more dynamic and better able to mold their shape when contacting cancer cells. AFM analysis revealed that c-Jun KO HEI-286 SCs were indeed stiffer than control HEI-286 SCs (Fig. 5 I,J). The difference of physical properties might originate from a re-arrangement of actin. We looked for actin organization in co-cultured HEI-286 SCs. Phalloidin staining revealed a difference of actin organization between control and c-Jun KO HEI-286 SCs. Actin was organized in stress fibers in control cells and as cortical actin in c-Jun KO cells (Fig. 5K). These findings suggest that SC c-Jun regulates actin organization and modifies cellular physical properties. In addition, c-Jun enables SCs to wrap cancer cells, conform to the microchannel, and create tracks supporting fast directional migration of cancer cells.

### c-Jun coordinates SC collective organization into TASTs that allow cancer invasion

We characterized the role of c-Jun in the 3D organization of SCs in Matrigel. The control HEI-286-SCs adopted an elongated shape and self-organized into linear, branched structures in the Matrigel, forming frequent end-to-end cellular contacts (Fig. 6A, left). In contrast, the c-Jun KO HEI-286 SCs adopted a more rounded shape (Fig. 6A, right), comparable to that of c-Jun KO murine SCs *in vivo* and *in vitro*(27) and formed disorganized clusters, demonstrating a fundamental role of c-Jun in the 3D self-organization of SCs into linear structures. No difference was observed in 2D between control and c-Jun KO SCs (Fig. 6A, inserts), showing the importance of the dimensional environment in SC organization. P-c-Jun was expressed in the 3D SC structures (Supplementary Fig. 6A). SCs treated with SP600125, a JNK inhibitor that blocks c-Jun phosphorylation (Supplementary Fig. 6B) also exhibited an inability to collectively organize into linear structures in 3D (Supplementary Fig. 6C). We next followed the organization of HEI-286 SCs using time-lapse microscopy and measured the maximum length of the SC structures formed from a single starting cell in 3D. After 80 hours and about 3–4 mitoses, the daughter cells of a control SC connected to create a branched, linear structure (Supplementary Video 7, left), that was longer than the structure created by c-Jun KO HEI-286 SCs (Fig. 6B, Sup Fig D) or SP600125 treated cells (Fig. 6C Sup Fig E). Tracking analysis of time-lapse movies indicated that the control daughter cells separated from one another and initially migrated in opposite directions before stretching out and oscillating back and forth to eventually connect with one another (Supplementary Fig. 6F, G). This sequence was a consistent initial step for replicating HEI-286 SCs to link together into 3D branched, linear structures. Single c-Jun KO HEI-286 SCs or SP600125-treated control SCs also underwent mitosis at least 3–4 times, but remained bunched together(Fig. 6C-F; Supplementary Videos 7 and 8). They exhibited defects in separation and migration away from each other after mitosis (Supplementary Fig. 6E,G; Supplementary Videos 7 and 8). To quantify daughter cell separation following mitosis, we performed time-lapse microscopy in microchannels. As in Matrigel, after cell division, daughter control HEI-286 SCs consistently separated and migrated away from each other (Supplementary Fig. 6H; Supplementary Video 9, top). They remained separated until the cells became more confluent in the microchannel. In contrast, daughter c-Jun KO HEI-286 SCs failed to migrate away from one another, and re-established contact after mitosis (Supplementary Fig. 6H; Supplementary Video 9, bottom). This was quantified by measuring the maximum distance of separation between two daughter HEI-286 SCs following mitosis (Supplementary Fig. 6I) and the percentage of daughter cells that separated from each other following mitosis (Supplementary Fig. 6J). These data provide insights into how cell separation is an integral component of HEI-286 SCs self-organization into 3D branched, linear structures. This process is driven by c-Jun and forms the basis for HEI-286 SC alignment into tracks.

**Fig. 6.**
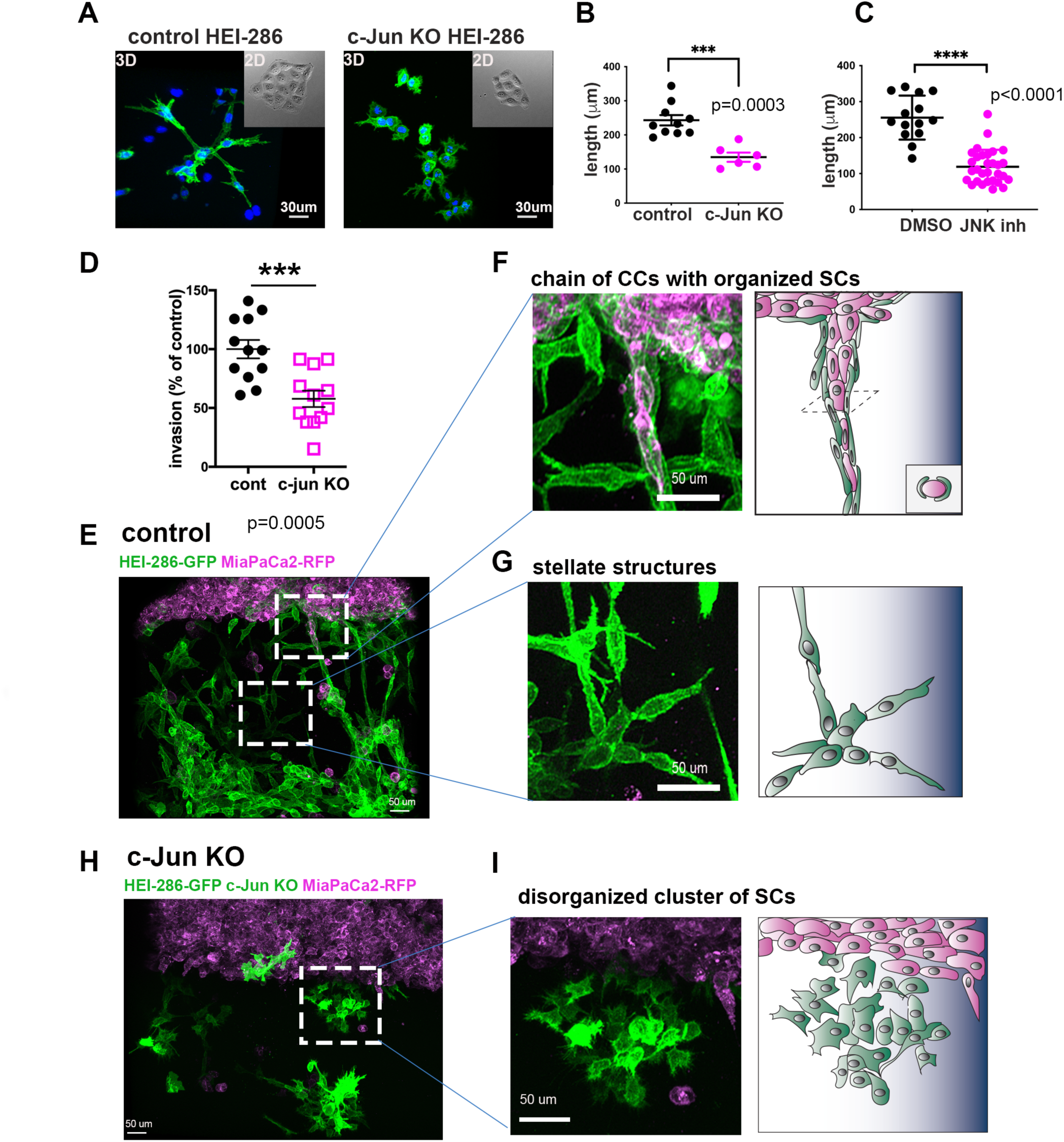
c-Jun coordinates SC collective organization and enhances cancer cell invasion in 3D Matrigel through track formation. **A**, Confocal images of control and c-Jun KO HEI- GFP-SCs in Matrigel showing a lack of SC organization in c-Jun KO HEI-286 SCs. Scale bars, 30 μm. **B**,**C** Quantification of the length of the SC structures created in Matrigel from one cell after 80h for control and c-Jun KO cells (control n=10, c-Jun KO n=6, mean ± SEM) (B) and JNKinhibitor treated cells (control n=14, JNKinh n=30, mean ± SEM) (C). **D**, Quantification of MiaPaCa2 invasion into a 3D Matrigel chamber in presence of control vs. c- Jun KO HEI-286 SCs grown in the Matrigel (n=12 measurements/condition, mean ± SEM). **E**, Maximum projection view of confocal images of MiaPaCa2 (magenta) invasion in presence of HEI-286-SCs (green) growing in Matrigel. Scale bar, 50 μm. **F**, Confocal image enlarged from (**E**) and schematic showing a chain of cancer cells aligned within a tubular, linear structure of control SCs in Matrigel. Scale bar, 50 μm. **G**, Confocal image enlarged from (**E**) and schematic showing branched structures created by aligned SCs in Matrigel. Scale bar, 50 μm. **H**, Maximum projection view of confocal images of MiaPaCa2 invasion in presence of c-Jun KO HEI-286-SCs growing in Matrigel. Scale bar, 50 μm. **I**, Confocal image enlarged from (**F**) and schematic showing a disorganized cluster of c-Jun KO HEI-286 SCs, lacking organization. Scale bar, 50 μm.

We assessed the role of SC c-Jun in the 3D model of cancer cell invasion in which we showed HEI-286 SCs organizing around MiaPaCa-2 cancer cells (Fig. 2P-R). Analysis of confocal images revealed that c-Jun KO HEI-286-SCs were less able to support cancer invasion than control HEI-286 SCs (Fig. 6D). As previously described, cancer cells were associated with the branched, linear structures of control HEI-286 SCs and aligned along these tracks extending into the Matrigel (Fig. 6E-G). In contrast, the c-Jun KO HEI-286 SCs failed to form linear structures but largely remained in disorganized clusters (Fig. 6H,I).

MiaPaCa-2 did not show affinity for the unstructured c-Jun KO HEI-286 SCs and failed to align as chains of cells, although they occasionally associated with individual c-Jun KO HEI- 286 SCs (Fig. 6H,I) These data demonstrate a fundamental role of c-Jun in the 3D self-organization of SCs into linear structures that serve as pathways for cancer cells.

### SC c-Jun promotes cancer cell neural invasion *in vivo*

To investigate the role of SC c-Jun in PNI *in vivo*, we injected Panc02 cancer cells into the sciatic nerves of c-Jun KO and WT mice to assess the resulting length of PNI proximally toward the spinal cord from the site of injection(35). Histological analysis of hematoxylin and eosin (H&E) staining of nerve longitudinal sections after 7 days showed a significant reduction of PNI length along the sciatic nerve in the c-Jun KO mice as compared with WT mice (Fig. 7A,B). Quantifying sciatic nerve function(36, 37) indicated that WT mice developed more severe nerve paralysis and hindlimb dysfunction 6, 10, and 14 days after injection as compared with the c-Jun KO SC mice (Fig. 7C-E).

**Fig. 7.**
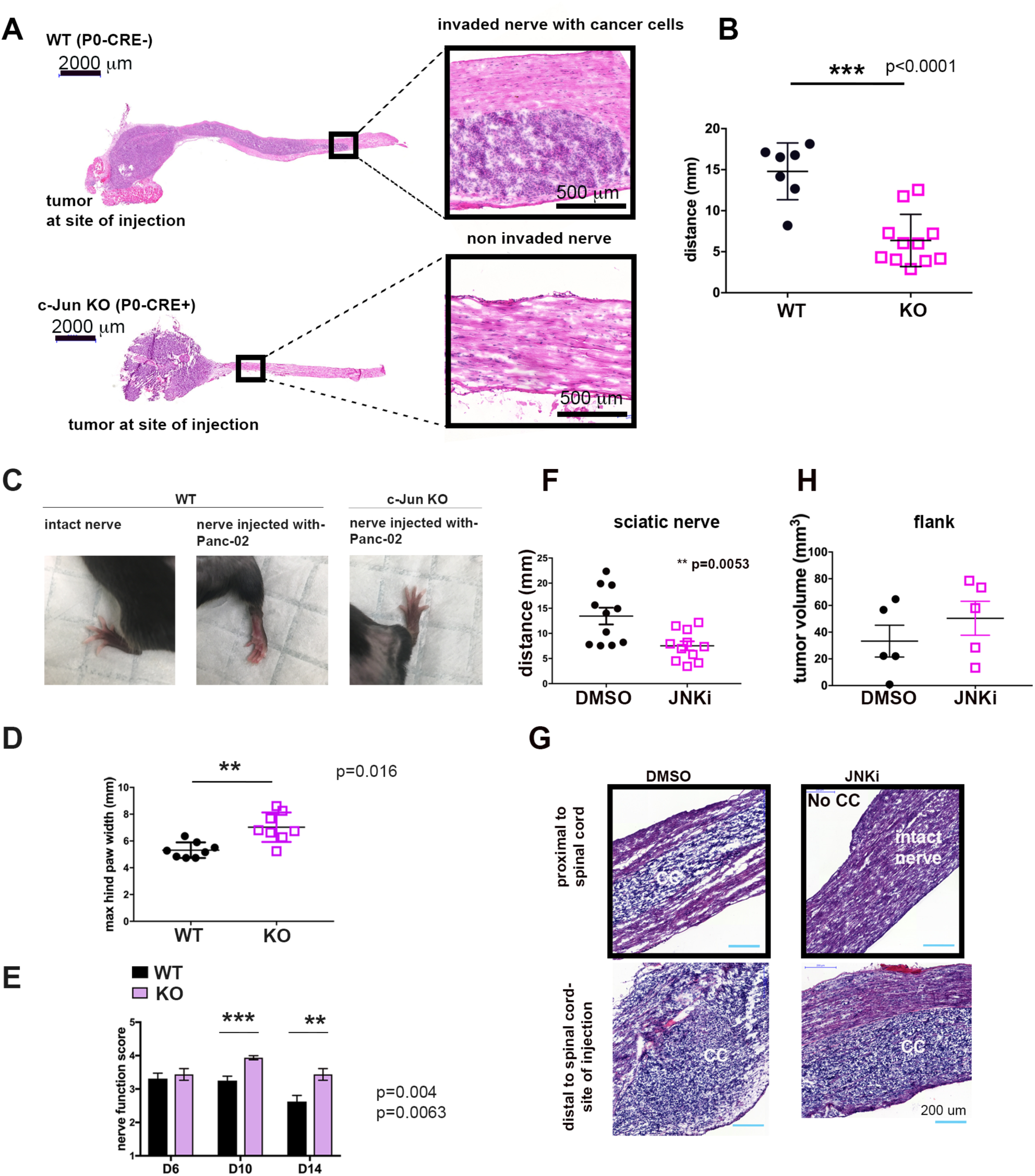
c-Jun reprogrammed SCs promote cancer invasion *in vivo.* **A**, Histological analysis of injected murine sciatic nerves in P0-CRE- (control) and P0-CRE+ (c-Jun KO) c- Jun^fl/fl^ mice. Representative samples of cancer invasion detected by H&E staining. Boxes show invasion of areas away from site of injection. Scale bars, 2000 μm. Box scale bars, 500 μm. **B**, Quantification of (E) (control n=7, c-Jun KO n=11, mean ± SEM). **C**, Representative images of murine hindlimbs after Panc02 cancer cell injections show less paralysis in c-Jun KO SC mice. **D**, Quantification of the maximum width of hind limb paw in P0-CRE- (control) vs. P0-CRE+ (c-Jun KO) c-Jun fl/fl mice 10 days after cancer injection. (n=8 mice/condition, mean ± SEM). **E**, Quantification of sciatic nerve function in P0-CRE- (control) vs. P0-CRE+ (c-Jun KO) cjun fl/fl mice (n=8 mice/condition, mean ± SEM). **F**, Effect of SP600125 on sciatic nerve invasion. Quantification of length of nerve invasion in mice injected with Panc02 and SP600125, or Panc02 and DMSO (n=11 mice/treatment, mean ± SEM). **G**, Representative H&E staining images of sciatic nerves injected with Panc02 and SP600125 vs. Panc02 and DMSO. Upper images are from proximal nerve regions at the spinal cord. No cancer is seen within the SP600125 injected nerve. Bottom images are from the site of injection, distal to spinal cord. CC is cancer cells. Scale bars, 200 μm. **H**, Effect of SP600125 on flank tumor growth. Quantification of tumor volume in mice co-injected with Panc02 and SP600125, and with Panc02 and DMSO. (n=5 mice/treatment, mean ± SEM).

To corroborate the role of SC c-Jun in PNI and to address the therapeutic utility of targeting c-Jun N-terminal Kinase (JNK), we applied the JNK inhibitor SP600125 to the same experiment described above using the murine sciatic nerve model. The addition of SP600125 significantly decreased the length of PNI caused by Panc02 cancer cells as compared with controls (Fig. 7F,G). SP600125 did not affect cancer growth or tumor volumes when Panc02 cells were injected into the murine flank (Fig. 7H), demonstrating that inhibitory effects of SP600125 were not due to a direct effect on Panc02 proliferation. Together, these data suggest that c-Jun activity in SCs can be effectively targeted by pharmacologically blocking JNK, and that this modulation of the nerve microenvironment can have therapeutic effects in inhibiting cancer progression along nerves.

## DISCUSSION

While the concept that cells from the TME contribute to cancer invasion is established, most reported mechanisms have focused on well-identified cells such as fibroblasts and immune cells. Our bioinformatics approach revealed the importance of non-myelinating Schwann cells, which are more difficult to detect than other cells from the TME using conventional tools. Our experiments demonstrate how these cells unconventionally lead to the invasive phenotype of the cancer cells. Our study describes a mechanism of invasion in which non-myelinating, activated SCs create Tumor Activated Schwann cell Tracks (TASTs), 3D structures that serve as a living scaffold to promote cancer invasion. However, this scaffold is not simply a passive structure. These SC tracks are composed of highly dynamic SCs that directly apply forces onto cancer cells to enhance their migration. These SCs exhibit a variety of complex behaviors that induce pushing, squeezing, and pulling forces on cancer cells. They are also highly pliable and allow cancer cells to pass alongside them in a confined space.

Our computational analysis of TCGA gene expression data uses a variety of different SC signatures. This work suggests a clinical importance of SCs in PDAC that has not been previously reported, revealing in particular the significance of gene profile associated with non-myelinating SCs. While the phenotype of myelinating cells is distinctive, SCs that do not make myelin such as non-myelinating (Remak) SCs and repair SCs in injured nerves share several molecular features. Our results indicate that cancer cells shift SC differentiation state from that of myelin SCs towards the gene expression characteristic of SCs that do not make myelin, including elevated expression of c-Jun and GFAP, a SC differentiation state that, in turn, promotes cancer cell migration and invasion. Unlike the myelinating SC signature, the non-myelinating SC signature significantly correlates with worse outcomes for patients in the dataset from TCGA. This is further supported by our newly defined cancer-exposed SC signature (HEI-mix) that reveals a strong correlation with worse PAAD survival. Our bulk TCGA gene expression analyses link the presence of these SCs with an invasive cancer phenotype and axonal guidance properties, suggesting that these SCs enable cancer invasion pathway activity while also utilizing SC axonal guidance programming.

Activated SCs have been shown to be present in a variety of cancers(18,38–41). We demonstrate that cancer cells activate SCs in a manner related to that seen after nerve injury and activate the transcription factor c-Jun. SCs organized in linear structures function to promote cancer invasion, recapitulating to some extent SC behavior following nerve injury. Following nerve transection, repair SCs migrate to traverse the gap, creating a linear bridge that links the cut ends together(29). SCs along the denervated distal nerve stump adopt an elongated and often branched morphology, enabling them to form regeneration tracks, Büngner bands, that guide re-growing axons(42). We show here that SCs adopt a similar phenotype in 3D gel, creating branching patterns, with elongated cells forming long tracks. In the cancer setting, these SC tracks enable opportunistic cancer invasion, allowing cancer cells to align and invade along TASTs extending into the 3D matrix. These findings suggest that cancer cells hijack the SC nerve repair program and are guided by SC tracks in a manner that is analogous to how regenerating axons are guided during nerve repair.

The identification of c-Jun as a key regulator of this cancer-associated SC phenotype may be harnessed for the treatment of SC-induced cancer invasion. We demonstrate that SC self-organization into TASTs, SC-mediated cancer cell migration and invasion, SC association with cancer cells *in vivo*, and SC-mediated increase in PNI *in vivo* are all attenuated with the loss of c-Jun activity. Importantly, the use of a JNK inhibitor was able to impede PNI in a mouse model, demonstrating that this mechanism can be therapeutically targeted.

## METHODS

### CONTACT FOR REAGENTS AND RESOURCE SHARING

Further information and requests for resources and reagents should be directed to and will be fulfilled by the Lead Contact Richard J. Wong.

### EXPERIMENTAL MODEL AND SUBJECT DETAILS

#### Mice

All mouse procedures were performed in accordance with institutional protocol guidelines at Memorial Sloan Kettering Cancer Center (MSK). C57BL/6J were obtained from the Jackson Laboratory (Stock No 000664). The protein myelin zero promoter specific Cre recombinase mice P0-CRE and P0-CRE cjun fl/fl were previously described (27). P0-CRE mT/mG were generated by crossing P0-CRE mice with mT/mG mice obtained from the Jackson Laboratory (Stock No 007676).

#### Cell lines and cell culture

The human non-neoplastic HEI-286-GFP stable cell lines and the pancreatic cancer cells MiaPaCa-2 RFP were previously described (18). The murine cell line Panc02 was from Dr. Min Li (43). c-Jun KO HEI-GFP cell lines were generated using the CRISPR-Cas9 technology. Two constructs were made at the MSK Gene Editing & Screening Core Facility using lentiCRISPRv2 vector (addgene 52961) and the following Jun targeting oligonucleotide sequences:

JUN-1F CACCGCCGTCCGAGAGCGGACCTTA

JUN-1R AAACTAAGGTCCGCTCTCGGACGGC

JUN-2F CACCGTCGGCGGCGCAGCCGGTCAA

JUN-2R AAACTTGACCGGCTGCGCCGCCGAC

DNA constructs were nucleofected in HEI-286-GFP using Amaxa (program T-020). A control cell line was generated using an empty vector. Stable cell lines were generated after selection with puromycin (3 μg/ml). We generated both polyclonal cell lines and a monoclonal cell line for the sequence number 2. Depletion of c-Jun in c-Jun K0-HEI was verified by Western blotting and immunofluorescence assays using standard protocols.

All cells were cultured in 5% CO2 at 37°C in Dulbecco’s modified Eagle’s medium (DMEM, Cellgro, Herndon, VA) containing 10% fetal bovine serum (Gemini, Woodland, CA) and 50 U/ml penicillin/streptomycin (Cellgro, Herndon, VA). Culture medium for the c-Jun KO HEI-GFP cells was supplemented with puromycin (3ug/ml). Culture medium for cells expressing GFP and RFP and Far red-PM670 were supplemented with G418 (50ug/ml). Cell lines were routinely screened to avoid mycoplasma contamination and maintained in a humidified chamber with 5% CO2 at 37°C.

#### Patient materials

All patients signed an approved informed consent before providing tissue samples. Patient samples were collected using a tissue-collection protocol approved by the MSK Institutional Review Board. The non-PDAC pancreatic specimens included one patient with a benign pancreatic cyst (with history of rhabdomyosarcoma) (1), two patients with well-differentiated pancreatic neuroendocrine tumors (2 and 3), and one patient with a solid pseudopapillary neoplasm (4).

### METHOD DETAILS

#### Murine sciatic nerve injection

Sciatic nerve injection procedures were carried as previously described (34, 35). Briefly, mice (P0-CRE c-Jun fl/fl, P0-CRE mT/mG and C57BL/6J mice) were anesthetized using isoflurane (1-3%), and their sciatic nerve exposed. Depending on the experiments, sterile PBS, murine pancreatic cancer cells Panc02 (fluorescent or not) (50 000 cells) were injected into the sciatic nerve under loop magnification using a 10 μl Hamilton syringe. For the assessment of SP600125 effect on PNI, Panc02 were injected in the nerves with SP600125 (50mM, ug/weight) or DMSO (control). After treatment with meloxicam for analgesia, the mouse was closed up with surgical sutures. Mice were followed for recovery every day for 72h and monitored for nerve function.

#### Murine sciatic nerve function

Sciatic nerve function was measured in control and c-Jun KO mice 6, 10 and 14 days after injection of Panc02 cells by using the nerve function score (36, 37). It was graded from 4 (normal) to 1 (total paw paralysis), according to hind limb paw response to manual extension of the body. The maximum hind paw width was measured at day 10 post-injection using a digital caliper.

#### Preparation of murine sciatic nerve sections

Seven days after injection, murine sciatic nerves were dissected up to the spinal cord and embedded in Tissue-Tek OCT. (Electron Microscopy Sciences). The specimen frozen blocks were serially sectioned longitudinally at respectively 5 μm thickness using Cryostat microtome (Leica CM1950). Sections were used for H&E and immunofluorescence staining. Morphological assessment and quantification of neural invasion by cancer cells in sciatic nerves were performed using Panoramic Viewer (3DHISTECH) on the H&E sections.

#### Immunofluorescence of murine sciatic nerve sections and human specimen sections

Murine frozen sections were fixed using 4% paraformaldehyde. Sections from human pancreatic specimens were obtained from paraffin blocks that were prepared using a standard protocol. Sections were permeabilized and blocked in 3% horse serum, 0.1% Triton X-100/ PBS for 1h. Primary antibodies (anti P-c-Jun 1:200, anti GFAP 1:1000, anti-cytokeratin 1:200) diluted in 0.1% horse serum, 0.1% Triton X-100/PBS were incubated O/N at 4°C. Sections were washed with PBS and detection was performed using an appropriate fluorescent secondary antibody (Alexa Fluor 488, 568, or 647, Invitrogen). Samples were mounted in DAPI containing antifade mounting medium. Slides were scanned using Flash Scanner (Perkin Elmer). Immunofluorescent sections were analyzed using Panoramic Viewer (3DHISTECH).

#### Immunofluorescence and actin staining of HEI-286 SCs co-cultured with MiaPaCa-2

GFP HEI-286 and RFP MiaPaCa2 (30,000 cells total, 1/1) were seeded on 35 mm glass bottom dish pre-coated with 20% Matrigel for 3 days. Cells were fixed in 4% paraformaldehyde/PBS for 15 minutes and washed 3 times with PBS. For immunofluorescence, cells were blocked in 3% horse serum, 0.1% Triton X-100/ PBS for 1h and incubated with antibody anti P-c-Jun diluted in 0.1% horse serum, 0.1% Triton X- 100/PBS O/N at 4°C. After washing, cells were incubated with Alexa Fluor 647 secondary antibody diluted in 0.1% horse serum, 0.1% Triton X-100/PBS containing DAPI. For actin staining, cells were incubated with Alexa Fluor Plus 647 Phalloidin (1:100) (#A2287, Invitrogen) for 30mins at room temperature and stained with DAPI for 30 minutes.

Images were taken using an inverted Axiovert 200M microscope (Zeiss) equipped with EC Plan-Neofluar 40X 0.3NA Ph1 lens and analyzed using Axiovision software and Fiji.

#### Invasion assay

Invasion assay was performed as previously described (18). Briefly, HEI-286 GFP or HEI- 286 c-Jun KO were seeded in 40 μl Matrigel matrix at 70 cells/μl in a two chamber insert (Ibidi GmbH, Germany) placed in glass-bottom 35mm MatTek dishes and grown for 6 days in DMEM 10% FBS. MiaPaCa-2 RFP (50,000 cells) were then added on top of the Matrigel.

After 3 days, samples were fixed in 4% paraformaldehyde for 30 min for imaging. The co-culture was imaged using a Leica SP5 or SP8 inverted confocal microscope with a 10X 0.4 NA lenses. (The z-stacks obtained with the 10X lens were used for quantification of invasion and were 1 mm thick with 5 μm step. Images were recorded at 12-bit resolution. Stacks were reconstructed in 3 dimensions (3D) using Imaris software (Bitplane) and red fluorescent structures corresponding to cancer cells were quantified using the Imaris software. The software determined the number of red cancer cells in an area of interest (size x=66, y=296, z=552 μm), that was about 300 μm below the surface of the Matrigel. The invasion was about 60 cells/area of interest and was considered 100%.)

Samples were turned upside down for imaging the interaction of HEI-GFP (control and c-Jun-KO) and MiaPaCa-2-RFP at the top of Matrigel. The lens used was 20X and the stack thickness was set for optimal resolution. At the top of the Matrigel, we counted 5.5 ± 1.73 chains of cancer cells surrounded by HEI-GFP per mm squared surface area with control HEI-GFP and about 4 disorganized KO HEI-FGP clusters per mm squared surface area.

#### Imaging SCs in 3D Matrigel

HEI-GFP (control and c-Jun KO) were mixed at 4°C with 100% Matrigel at a concentration of 70 cells/μl. Drops of 10μl drops of the Matrigel containing SCs were placed in glass bottom dishes. The samples were then incubated at 37°C for 30 min to allow polymerization. Culture medium was added and the samples were incubated at 37°C 5% CO2 until imaging. In some experiments, 10μM SP600125 or DMSO was added to the medium at the time of plating. Images of cell clusters were made 5 days after seeding using Leica confocal microscope with a 20X 0.7 NA lenses. Time-lapse imaging was performed 24 hours after seeding at the 2-cell stage using a motorized stage inverted Axiovert 200M microscope (Zeiss) equipped with EC Plan-Neofluar 10X 0.3NA Ph1 lens and temperature and CO2 controllers. Images were recorded every 10 min for 72h.

#### Imaging microchannels

Microchannels (10-20μm width, 10μm height) (4dcell, France) were coated with fibronectin (10μg/ml) (Sigma-Aldrich F1141) for 1h at RT and washed 3 times with PBS before incubating with the cell culture medium for 1 hours at 37°C and 5%CO^2^. A total of 10 000 cells were placed in the wells. When SCs were combined with cancer cells, 5000 of each cell types were placed in the wells that connect the channels. Cells were recorded 24 hour or at indicated time after loading. Migrating cells were recorded overnight with inverted microscope (Zeiss) at 37°C with 5% CO^2^ atmosphere and a 10X objective (0.3NA) at one image every 10 min for 24h. For some experiments, still confocal images and time-lapse images were taken using inverted confocal microscope (Leica, SP5). Time-lapse images were taken for 12 h every 10 min.

#### Atomic force microscopy

Glass bottom petri dishes (FluoroDish FD5040, World Precision Instruments) containing HEI-286 SCs with MiaPaCa-2 cancer cells were used for the acquisition of stiffness maps. Experiments were performed at 37°C with a MFP-3D-BIO AFM microscope (Oxford Instruments). Fluorescent images of GFP-expressing HEI-286 SCs and RFP-expressing MiaPaCa-2 cancer cells were acquired together with the stiffness maps, using the inverted optical microscope (Zeiss AxioObserver Z1) integrated with the AFM microscope. Cantilevers with colloidal borosilicate probes were used for experiments, having diameter of 5 μm and nominal spring constant k = 0.1 N/m (Novascan). Before each experiment, the exact cantilever spring constant was determined with the thermal noise method and the optical sensitivity was determined using a glass bottom petri dish filled with PBS as an infinitely stiff substrate. Stiffness maps of 80x80 μm^2^ (20x20 points) were collected in areas containing the cells and the substrate, used as reference, at the rate of 1.5 Hz for a complete single approach/withdraw cycle. A trigger point of 1 nN was used to ensure maximum sample penetration of less than 1 micron. Force curves in each map were fitted according to the Hertz model using the routine implemented in the MFP3D AFM (Igor Pro, WaveMetrics). Data fitting was performed in the range from 0–60% of the maximum applied force, by setting tip Poisson ν_tip_ = 0.19, tip Young’s modulus E_tip_ = 68 GPa and sample Poisson ν_sample_ = 0.45 (44). To account for possible substrate effects, a threshold of 500 nm was chosen for the selection of the stiffness data, from the corresponding topographical maps of each cell collected *in situ* with the stiffness maps, to account for possible substrate effects. Extraction of the stiffness values from the raw Igor Binary Wave (.ibw) data within the mask region was obtained by means of a home-built routine implemented in Igor (Igor Pro, WaveMetrics) and a custom script in MATLAB (MathWorks). Data visualization was performed using OriginPro (OriginLab) and GraphPad Prism. Stiffness analysis was performed only on SCs physically connected to cancer cells, as inferred from the respective fluorescence signal.

#### Cell sorting, RNAseq library preparation and analysis

RNASeq data were from three independent biological replicates. Four million GFP- expressing HEI-286 cells were grown alone or mixed with RFP-expressing MiaPaCa-2 for four days (1:1). Cells were trypsinized, pelleted, re-suspended in PBS and passed through a 35μm cell strainer to yield single cells. Live (DAPI negative) GFP cells were sorted on a BD SORP FACSAria^TM^ IIu equipped with a 488 nm laser and a 525/50 nm bandpass filter to excite and detect GFP, a 561 nm laser and a 610/20 nm bandpass filter to excite and detect mRFP, and a 405 nm laser and 450/40 nm bandpass filter to excite and detect DAPI.

RNA was extracted from cells by using RNeasy mini kit (QIAGEN, cat #74104) as per the manufacturer’s instructions. RNAseq was performed by the MSK Integrated Genomics Operation. After RiboGreen quantification and quality control by Agilent BioAnalyzer, 500ng of total RNA underwent polyA selection and TruSeq library preparation according to instructions provided by Illumina (TruSeq Stranded mRNA LT Kit, catalog # RS-122-2102), with 8 cycles of PCR. Samples were barcoded and run on a HiSeq 2500 in a 50bp/50bp paired end run, using the HiSeq SBS Kit v4 (Illumina). An average of 58 million paired reads was generated per sample. At the most the ribosomal reads represented 5% of the total reads generated and the percent of mRNA bases averaged 79%. Sequence data processing was performed by the Bioinformatics Core at MSK. Output data (FASTQ files) were mapped to the human genome using the rnaStar aligner that mapped reads genomically and resolved reads across splice junctions. Output SAM files were post processed using PICARD tools and converted into BAM format. The expression count matrix from the mapped reads was computed using HTSeq (www-huber.embl.de/users/anders/HTSeq), and the raw count matrix was processed using the R/Bioconductor package DESeq (www.huber.embl.de/users/anders/DESeq) to normalize the full dataset and analyze differential expression between sample groups. 1092 out of 20245 genes were found differentially regulated in the co-cultured HEI-286 SCs as compared with HEI-286 SCs alone (fold change>2, p<0.05). Of those, 878 were upregulated. Because SCs are phagocytic and internalized material from cancer cells, consistent with its phagocytic role in Wallerian degeneration, genes found highly expressed in cancer cells were filtered out from the list of upregulated genes in SCs grown in presence of cancer cells using microarray data of HEI-286 and MiaPaCa-2 cells. The RNA of these HEI-286 and MiaPaCa-2 cells were extracted, cDNA were generated and applied to the Illumina HumanHT12 v4. 697 genes remained upregulated and were used for Gene Ontology (GO) pathway analysis using the KEGG 2019 human database. Genes found highly expressed in cocultured HEI-286 compared with HEI-286 grown alone, and cocultured c-Jun-KO HEI-286 compared with c-Jun-KO-HEI-286 grown alone, were submitted to GO enrichment analysis using a web-based tool (https://amp.pharm.mssm.edu/Enrichr/) for pathway analysis using the KEGG 2019 dataset. Heatmaps were generated using Morpheus.

### QUANTIFICATION AND STATISTICAL ANALYSIS

#### Bioinformatics analysis of correlations between SC gene signature scores or JUN expression and clinical outcomes in TCGA patients

Signatures for SCs were obtained from Tabula-Sapiens using OnClass(30) (http://tabula-sapiens-onclass.ds.czbiohub.org/), from PanglaoDB (45)(https://panglaodb.se/markers.html), and from a single cell analysis study of the pancreas(46). The activities of SC and other pathways were compared across tumor samples using a newly developed IPAS scores (31) (https://calina01.u.hpc.mssm.edu/pathway_assessor/). TCGA clinical data, including survival and new tumor events (NTE), were downloaded from the data portal of the Broad Institute (https://gdac.broadinstitute.org). Cox proportional hazards regression analysis was performed to examine the correlation between SC signature IPAS scores and patient outcomes (survival and NTE). The Survival package of R.3.6.0 was utilized to calculate log-rank P values. The optimal patient separation by IPAS score that yielded the lowest p value in clinical outcome was selected to separate the patients into high versus low score groups as an exploratory approach. The survival differences were visualized by generating Kaplan–Meier survival plots using Prism 9.

#### Quantification of cancer cells invasion in 3D assay

The z-stacks obtained with a 10X lens were used for quantification of invasion and were 1 mm thick with 5μm step. Images were recorded at 12-bit resolution. Stacks were reconstructed in 3 dimensions (3D) using Imaris software (Bitplane) and red fluorescent structures corresponding to cancer cells were quantified using the Imaris software. The software determined the number of red cancer cells in an area of interest (size x=66, y=296, z=552 μm), that was about 300μm below the surface of the Matrigel. The invasion was about 60 cells/area of interest and was considered 100%.

#### Quantification of length of SC structures in Matrigel, distance after division, SC separation after division and cancer cell passing a SC

Axiovision software (Zeiss) was used to measure the length of the structures created in Matrigel from two SCs after 72h, to measure the distance between two SCs in microchannels after division, to quantify cell separation after division in microchannels, and to quantify the number of events, cancer cells passing a SC in microchannels. To measure the distance between two SCs after division in microchannels, we selected dividing cells that had sufficient space to migrate away from each other. Distance measurement was done within 400 min after the division and included the space between the cells and the size of the cells. To quantify SC separation after division, dividing cells that had sufficient space to migrate away from each other were selected and counted as positive if the cells were separated 400 min after division. To quantify the number of events, cancer cells passing a SC in microchannels, single moving cancer cells that encountered a SC were selected. They were counted as positive if the cancer cell went through the SC. Analyses were performed visually.

#### Quantification of SC wrapping

Confocal xz projection images of SCs and cancer cells in contact within microchannels were analyzed to quantify SC wrapping around cancer cells. SCs were considered as wrapping cancer cells if they covered more than 60% of the cancer cell.

#### Quantification of instantaneous velocity

PIV analysis was performed by PIVlab version 2.36 (Time-Resolved Digital Particle Image Velocimetry Tool for MATLAB, developed by W. Thielicke and E. J. Stamhuis, http:// pivlab.blogspot.com). Images from time-lapse movies were first doubled in size and save as bmp files in Fiji. Using PIV, the cells were selected for every individual frame. Images were calibrated and analyzed.

#### Statistical analysis

Pairwise comparisons were conducted using unpaired two-tailed Student’s t-test. Statistical significance was defined at p values less than 0.05. Statistical analyses were performed using Prism 7 (GraphPad Software, Inc.).

## DATA AND SOFTWARE AVAILABILITY

The accession numbers for the RNA-seq and microarray data reported in this paper are GSE180710 and GSE180971, respectively.

## AUTHOR CONTRIBUTIONS

S.D. and R.W. designed, analyzed data and wrote the manuscript. S.D., L.G., A.P., Y.Y., C.H.C., A.L. and E.K. performed experiments. L.T. and E.V. analyzed the human histological sections. A.C. performed atomic force microscopy experiments and analyzed the data. T.O. performed analyzis of imaging experiments. B.R. performed bionformatic analysis. K.J. edited the manuscript and provided c-Jun KO mice. All authors read, pre-edited and approved the manuscript.

## ACKNOWLEDGMENTS

We acknowledge the technical services provided by MSK core facilities: the Molecular Cytology and Flow Cytometry facilities, the Integrated Genomics Operation Core, the Bioinformatics Core, and the Animal Imaging Core. We thank Ning Fan for assistance with sectioning specimens. The Integrated Genomics Operation Core was funded by the NCI Cancer Center Support Grant (CCSG, P30 CA08748), Cycle for Survival, and the Marie-Josée and Henry R. Kravis Center for Molecular Oncology. This work was supported by NIH CA 219534 (RW) and NIH P30 CA008748 (MSK CCSG).

## SUPPLEMENTARY FIGURE LEGENDS

**Supplementary Fig. 1 | Correlation of SC signatures and overall survival in TCGA patients with PAAD. A-G**, Kaplan–Meier curves of overall survival (OS) and new tumor event (NTE) with high or low scores for different signatures of SC in 178 TCGA PAAD patients. SC signature from Tabula Sapiens (**A**); SC signature from PanglaoDB (**B**); signature from Tosti et al. (46)(**C**); Terminal SC from Tabula Sapiens (**D**); Glial cell signature from Tabula Sapiens (**E**); SC precursor from Tabula Sapiens (**F**); Immature SC from Tabula Sapiens (**G**). **H–J**, Heatmaps of cell signatures correlating with scores for myelinating SC signature (**H**), non-myelinating SC signature (**I**) and HEImix signature (**J**) in TCGA PAAD patients.

Supplementary Fig. 2 | Cancer cells are closely associated with SCs in PDAC specimens and do not invade in absence of SCs in a 3D assay. A, Examples of uneven (top) and even (bottom) distribution of GFAP (green) staining in nerves of PDAC specimens. Nerves without (left) and with (right) visible cancer cells. Middle images are H&E of adjacent sections of right images. Ratios (bottom right corner) indicate number of specimens with even GFAP distribution over the total number of specimens. Scale bars 100 μm. B, GFAP+ SC (green) wrapping cytokeratin (CK) expressing cancer cells (magenta) in PDAC specimens. C-D, Absence of chain of cancer cells in absence of SCs or with NIH 3T3 fibroblasts. Scale bar, 50 μm. C, Confocal images of MiaPaCa2-RFP cells seeded on top of a Matrigel chamber and imaged after 6 days. Scale bar, 150 μm. D, Confocal images of MiaPaCa2-RFP cells taken 6 days after adding them on top of a Matrigel chamber previously seeded with NIH 3T3-GFP fibroblasts for 5 days. Scale bars 150 and 50 μm.

**Supplementary Fig. 3 | HEI-286-GFP SCs and MiaPaCa-2-RFP cancer cell microchannel assay. A**, Confocal images of HEI-GFP and MiaPaCa2-RFP within microchannels in longitudinal (xy) and transverse sections (xz) showing SCs fully occupying the channel (xz 2) or either wrapping partially (xz 3) or completely (xz 1) a cancer cell. Scale bars, 50 μm. **B**, Schematic showing the microchannel assay sequence. (1) HEI-286 SCs are first seeded in wells and enter microchannels. (2) MiaPaCa-2 cancer cells are seeded in the wells and (3) enter the microchannels occupied with HEI-286 SCs. **C**, Fluorescent images of time-lapse movie showing HEI-286 SCs (green) squeezing a cancer cell (magenta). Blue arrow indicates HEI-286 SC movement and white arrow indicates cancer cell displacement. Time is h:min. Scale bar, 20 μm.

**Supplementary Fig. 4 | c-Jun and P-c-Jun in SCs. A**, Western-Blot of HEI-286 SCs showing loss of c-Jun expression using either construct. **B**, Immunofluorescence of c-Jun in HEI-286 SCs showed diminished nuclear c-Jun expression with either construct. **C**, Proliferation of control and c-Jun KO HEI-286-SCs. **D**, Heatmaps of expression of axon guidance genes that are upregulated in co-cultured HEI-286 cells (HEI mix) as compared to HEI-286 grown alone (HEI). Each value is the mean of three biological replicates. **E**, Gene set enrichment analysis (GSEA) assessing upregulated c-Jun nerve-repair genes (27) in co-cultured HEI-286 compared to HEI-286 SCs. NES is normalized enrichment score. **F**, Inferred Pathway Activation/Suppression (IPAS) scores for the murine c-Jun nerve repair genes signature (27). **G**, GSEA assessing upregulated c-Jun genes in co-culture HEI versus grown alone HEI-286 in cut sciatic nerves compared to uncut sciatic nerves (27). NES is normalized enrichment score.

**Supplementary Fig. 5 | SC c-Jun supports both SC and cancer cell migration. A**, Quantification of control and c-Jun KO HEI-286-SC speed in two dimensions. **B**, Quantification of control and c-Jun KO HEI-286-SC speed in three dimensions in microchannels of sizes from 10 to 20 μm. **C**, Quantification of control and c-Jun KO HEI- 286-SC speed in three-dimension. **D**, Quantification of length of columns formed by MiaPaCa-2 combined with WT HEI-286 SCs or with c-Jun KO HEI-286-SCs. **E**, Individual tracks of cancer cells in microchannels occupied by control vs. c-Jun KO HEI-286 SCs. (n=21 in each group).

**Supplementary Fig. 6 | WT and c-Jun KO SCs behavior. A**, Confocal images of phospho-c-Jun (P-c-Jun) staining in HEI-286 GFP SCs in Matrigel. **B**, Effect of SP600125 on P-c-Jun expression in HEI-286 GFP SCs. **C**, Confocal images of HEI-GFP-SCs treated with JNKi SP600125 or DMSO in Matrigel showing lack of SC organization in JNKi-treated cells. Scale bars, 30 μm. **D-E**, Images of cells from 72h time-lapse movies of control and c-Jun KO HEI- 286 SCs (D) and DMSO or JNK inhibitor treated HEI 286 SCs (E) grown in 3D Matrigel with colored cell tracking. Scale bar, 60 μm. **F**, Images of WT SCs from time-lapse movies showing formation of a branched organized structure in 3D: cells separate after division (1); cells stretch and reconnect, reestablishing contacts (2); cells divide, organize in a structure that spreads out (3). Time in minutes. Scale bar, 60 μm. **G**, Schematic show the separation and positioning of two daughter HEI-286 SCs after mitosis in Matrigel is c-Jun-dependent. **H**, Time-lapse images of a dividing control and c-Jun-KO HEI-286 SCs and the two daughter cells in microchannels. **I**, Quantification of (**H**), the distance between two daughter cells following mitosis for control vs. c-Jun KO HEI-286 SCs in a 400 min time period (control n=16, c-Jun KO n=12, mean ± SEM). **J**, Quantification of percentage of daughter cells that separate following mitosis for control versus c-Jun KO HEI-286 SCs. (4 independent experiments, n>20 cells per condition in each experiment, mean ± SEM)

## SUPPLEMENTARY VIDEO LEGENDS

**Supplementary Video 1. MiaPaCa2 cancer cell (magenta) enter and migrate within a microchannel occupied by HEI-286 SCs (green).** Maximum projection of confocal images showing cancer cell (top), SCs (middle) and both cancer cell and SCs (bottom). Time lapse with images taken every 10 min. Scale bar, 30 μm. See Fig. 3C.

Supplementary Video 2. MiaPaCa2 cancer cells (magenta) migrate within a microchannel while being wrapped by HEI-286 SCs (green). Maximum projection of confocal images showing cancer cells (top), SCs (middle) and both cancer cells and SCs (bottom). Time lapse with images taken every 10 min. Scale bar, 30 μm. See Fig. 3D.

Supplementary Video 3. An HEI-286 SC (green) pushes a MiaPaCa2 cancer cell (magenta). Time lapse with images taken every 10 min. Scale bar, 50 μm. See Fig. 3E.

**Supplementary Video 4. HEI-286 SCs (green) squeeze a MiaPaCa2 cancer cell (magenta) which then moves to the left.** Maximum projection of confocal images showing both SCs and cancer cell (top), only cancer cell (middle) and only SCs (bottom). Time lapse with images taken every 10 min. Scale bar, 40 μm. See Supplementary Fig. 3C.

**Supplementary Video 5. HEI-286 SCs (green) pull a MiaPaCa2 cancer cell (magenta).** Time lapse with images taken every 10 min. Scale bar, 50 μm. See Supplementary Fig. 3H.

Supplementary Video 6. Time-lapse movie showing the behavior of a cancer cell (magenta) after contact with HEI-286 SCs (green) contact in a microchannel. The cancer cell passes next to a conntrol HEI-286 SC (top), but is blocked by a c-Jun KO HEI-286 SC. Images were taken every 10 min. Scale bars, 50 μm. See Fig. 5G.

Supplementary Video 7. Time-lapse movie of control (left) and c-Jun KO (right) SCs showing formation of an organized structure in control SCs, but not in c-Jun KO-SCs. Images were taken every 10 min. Scale bars, 50 μm. See Supplementary Fig. 6D.

**Supplementary Video 8. Time-lapse movie of SCs treated with JNKi SP600125 (right) or DMSO (left).** Unlike the JNKi-treated SCs, the DMSO-treated SCs organize into organized structures. Images were taken every 10 min. Scale bars, 50 μm. See Supplementary Fig. 6E.

Supplementary Video 9. Time-lapse images of a dividing control SC (top) and c-Jun-KO SC (bottom) and their daughter cells in microchannels. Images were taken every 10 min. Scale bars, 50 μm. See Supplementary Fig. 6H.

