## Supplementary material for "Reprogrammed Schwann Cells Organize into Dynamic Tracks that Promote Pancreatic Cancer Invasion": Sup Fig1

### Sup Fig 1

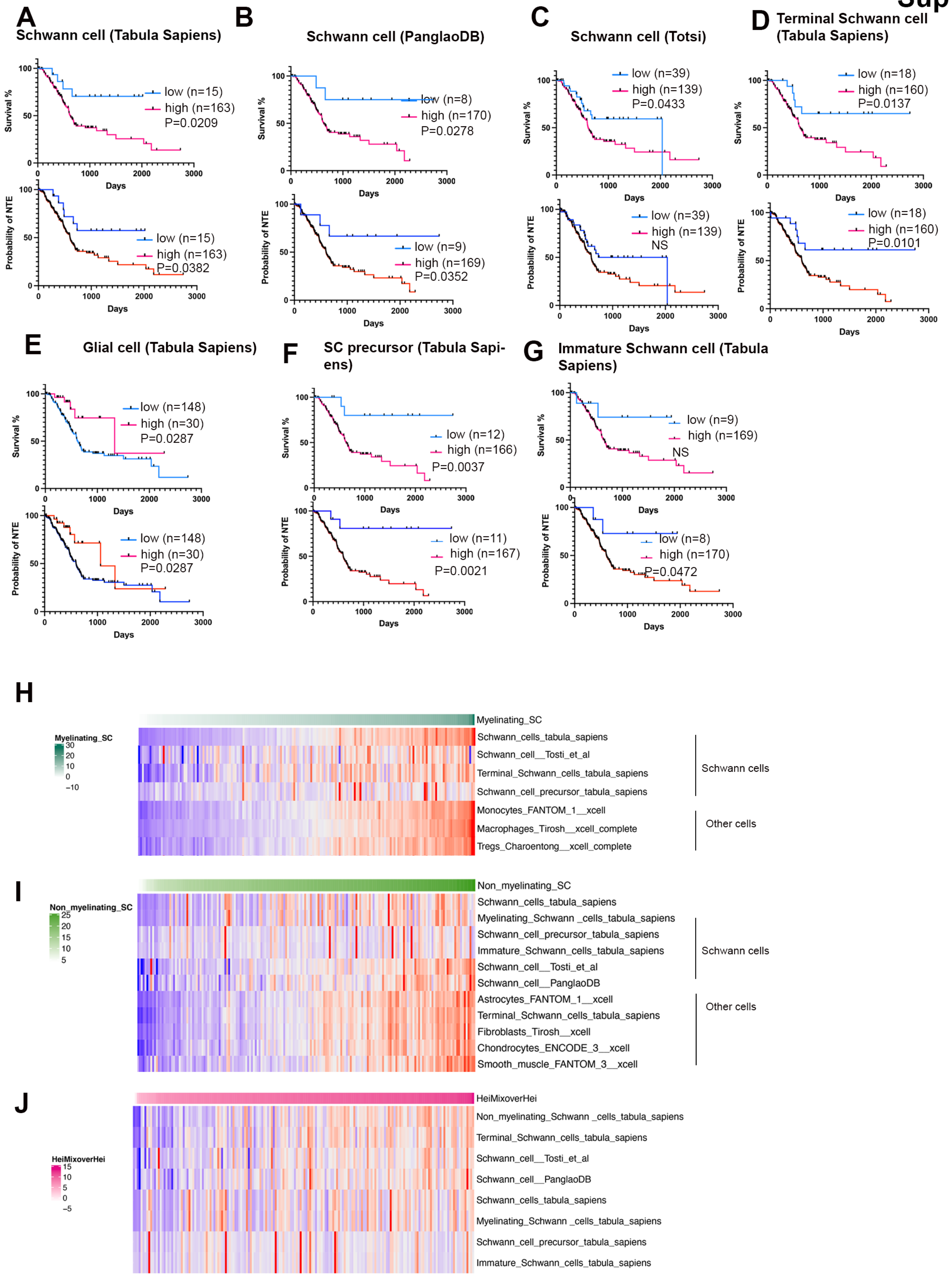


**Supplementary Fig. 1 | Correlation of SC signatures and overall survival in TCGA patients with PAAD.** **A-G**, Kaplan–Meier curves of overall survival (OS) and new tumor event (NTE) with high or low scores for different signatures of SC in 178 TCGA PAAD patients. SC signature from Tabula Sapiens (A); SC signature from PanglaoDB (B); signature from Tosti et al. (47)(C); Terminal SC from Tabula Sapiens (D); Glial cell signature from Tabula Sapiens (E); SC precursor from Tabula Sapiens (F); Immature SC from Tabula Sapiens (G). **H–J**, Heatmaps of cell signatures correlating with scores for myelinating SC signature (H), non-myelinating SC signature (I) and HEImix signature (J) in TCGA PAAD patients.
