## Supplementary material for "Reprogrammed Schwann Cells Organize into Dynamic Tracks that Promote Pancreatic Cancer Invasion": Sup Fig2

Sup Fig 2

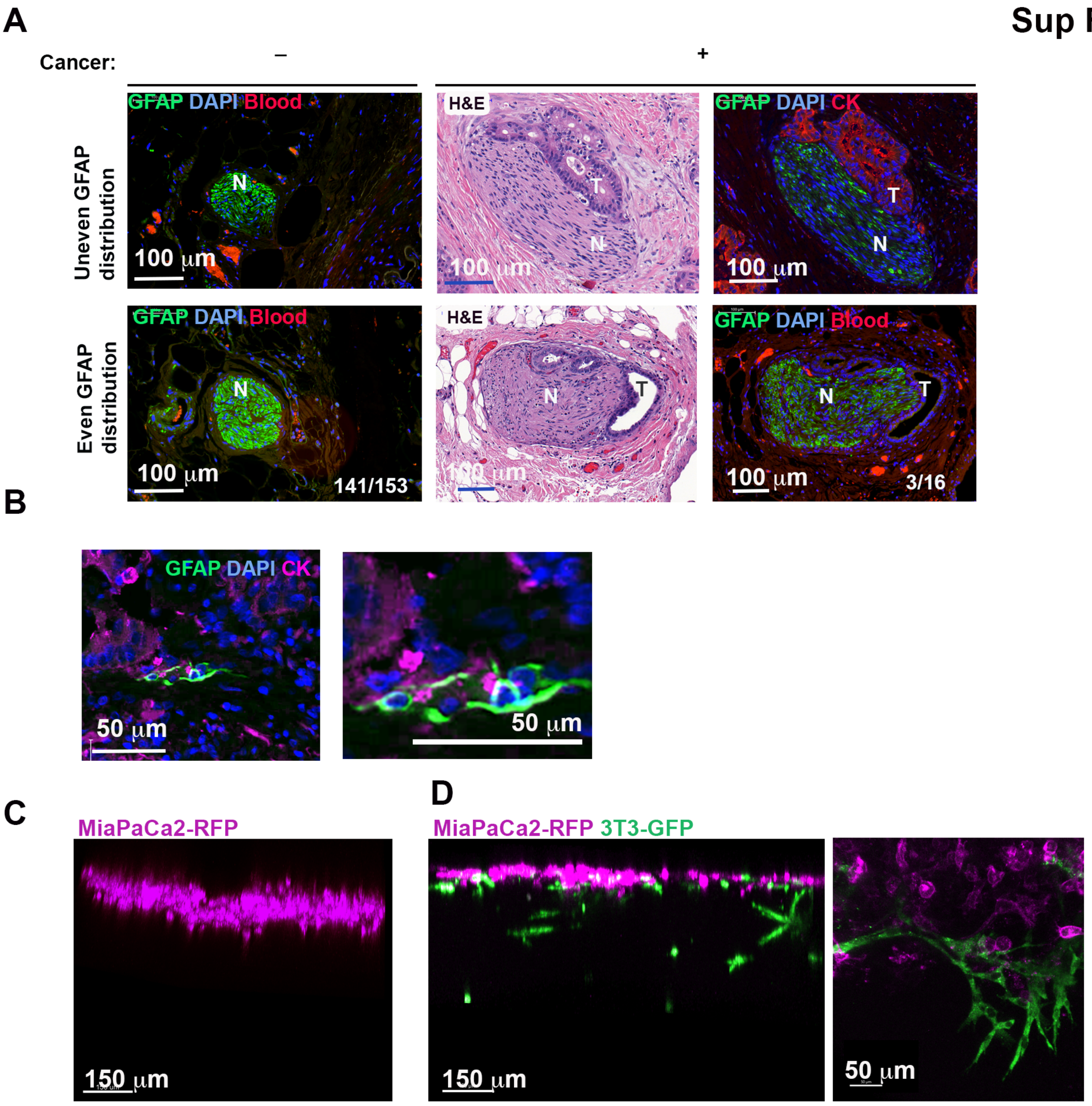


**Supplementary Fig. 2 |Cancer cells are closely associated with SCs in PDAC specimens and do not invade in absence of SCs in a 3D assay**. **A**, Examples of uneven (top) and even (bottom) distribution of GFAP (green) staining in nerves of PDAC specimens. Nerves without (left) and with (right) visible cancer cells. Middle images are H&E of adjacent sections of right images. Ratios (bottom right corner) indicate number of specimens with even GFAP distribution over the total number of specimens **B**, GFAP+ SC (green) wrapping cytokeratin (CK) expressing cancer cells (magenta) in PDAC specimen **C**-**D** Absence of chain of cancer cells in absence of SCs or with NIH 3T3 fibroblasts. **C,** Confocal images of MiaPaCa2-RFP cells seeded on top of a Matrigel chamber and imaged after 6 days. **D**, Confocal images of MiaPaCa2-RFP cells taken 6 days after adding them on top of a Matrigel chamber previously seeded with NIH 3T3-GFP fibroblasts for 5 days.
