## Supplementary material for "Reprogrammed Schwann Cells Organize into Dynamic Tracks that Promote Pancreatic Cancer Invasion": Sup Fig3

### Sup Fig 3

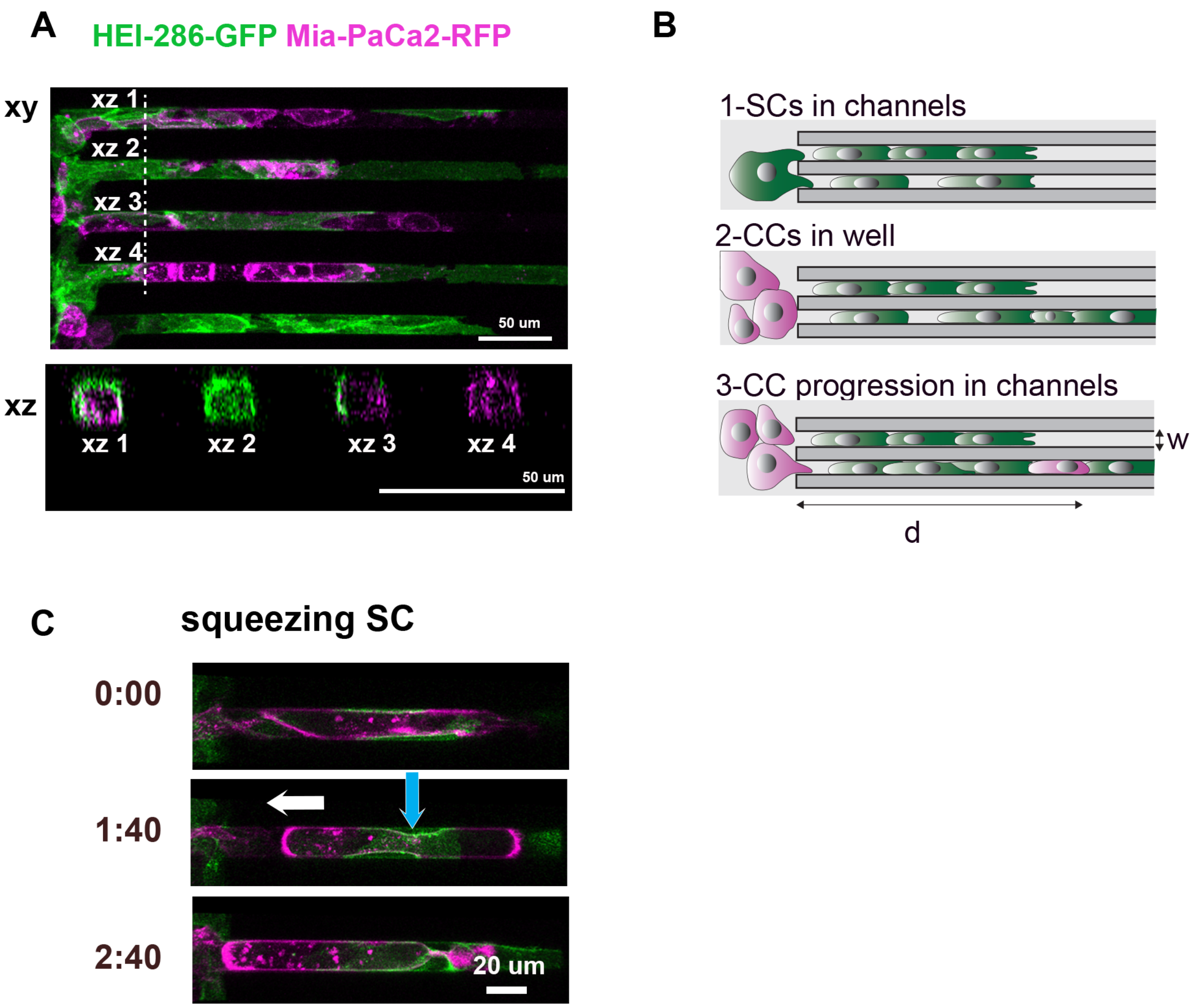


**Supplementary Fig. 3 | HEI-286-GFP SCs and MiaPaCa-2-RFP cancer cell microchannel assay. A,** Confocal images of HEI-GFP and MiaPaCa2-RFP within microchannels in longitudinal (xy) and transverse sections (xz) showing SCs fully occupying the channel (xz 2) or either wrapping partially (xz 3) or completely (xz 1) a cancer cell. **B**, Schematic showing the microchannel assay sequence. (1) HEI-286 SCs are first seeded in wells and enter microchannels. (2) MiaPaCa-2 cancer cells are seeded in the wells and (3) enter the microchannels occupied with HEI-286 SCs. **C**, Fluorescent images of time-lapse movie showing HEI-286 SCs (green) squeezing a cancer cell (magenta). Blue arrow indicates HEI-286 SC movement and white arrow indicates cancer cell displacement. Time is h:min.
