## Supplementary material for "Reprogrammed Schwann Cells Organize into Dynamic Tracks that Promote Pancreatic Cancer Invasion": Sup Fig4

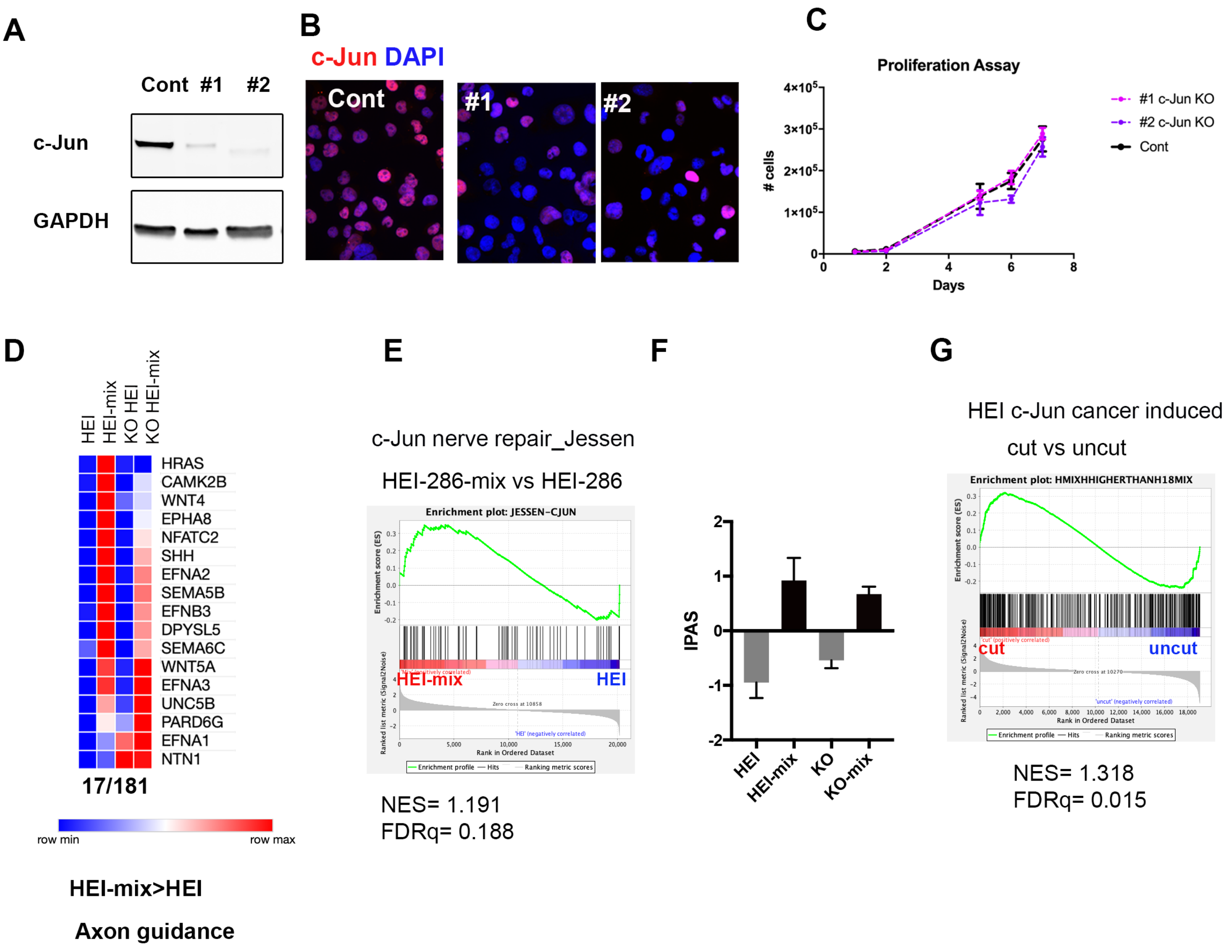


**Supplementary Fig. 4 | c-Jun and P-c-Jun in SCs**. **A**, Western-Blot of HEI-286 SCs showing loss of c-Jun expression using either construct. **B**, Immunofluorescence of c-Jun in HEI-286 SCs showed diminished nuclear c-Jun expression with either construct. **C**, Proliferation of control and c-Jun KO HEI-286-SCs. **D**, Heatmaps of expression of axon guidance genes that are upregulated in co-cultured HEI-286 cells (HEI mix) as compared to HEI-286 grown alone (HEI). Each value is the mean of three biological replicates. **E**, Gene set enrichment analysis (GSEA) assessing upregulated c-Jun nerve-repair genes (27) in co-cultured HEI-286 compared to HEI-286 SCs. NES is normalized enrichment score. **F**, Inferred Pathway Activation/Suppression (IPAS) scores for the murine c-Jun nerve repair genes signature (27). **G**, GSEA assessing upregulated c-Jun genes in co-culture HEI versus grown alone HEI-286 in cut sciatic nerves compared to uncut sciatic nerves (27). NES is normalized enrichment score.
