## Supplementary material for "Reprogrammed Schwann Cells Organize into Dynamic Tracks that Promote Pancreatic Cancer Invasion": Sup Fig5

**Sup Fig 5**

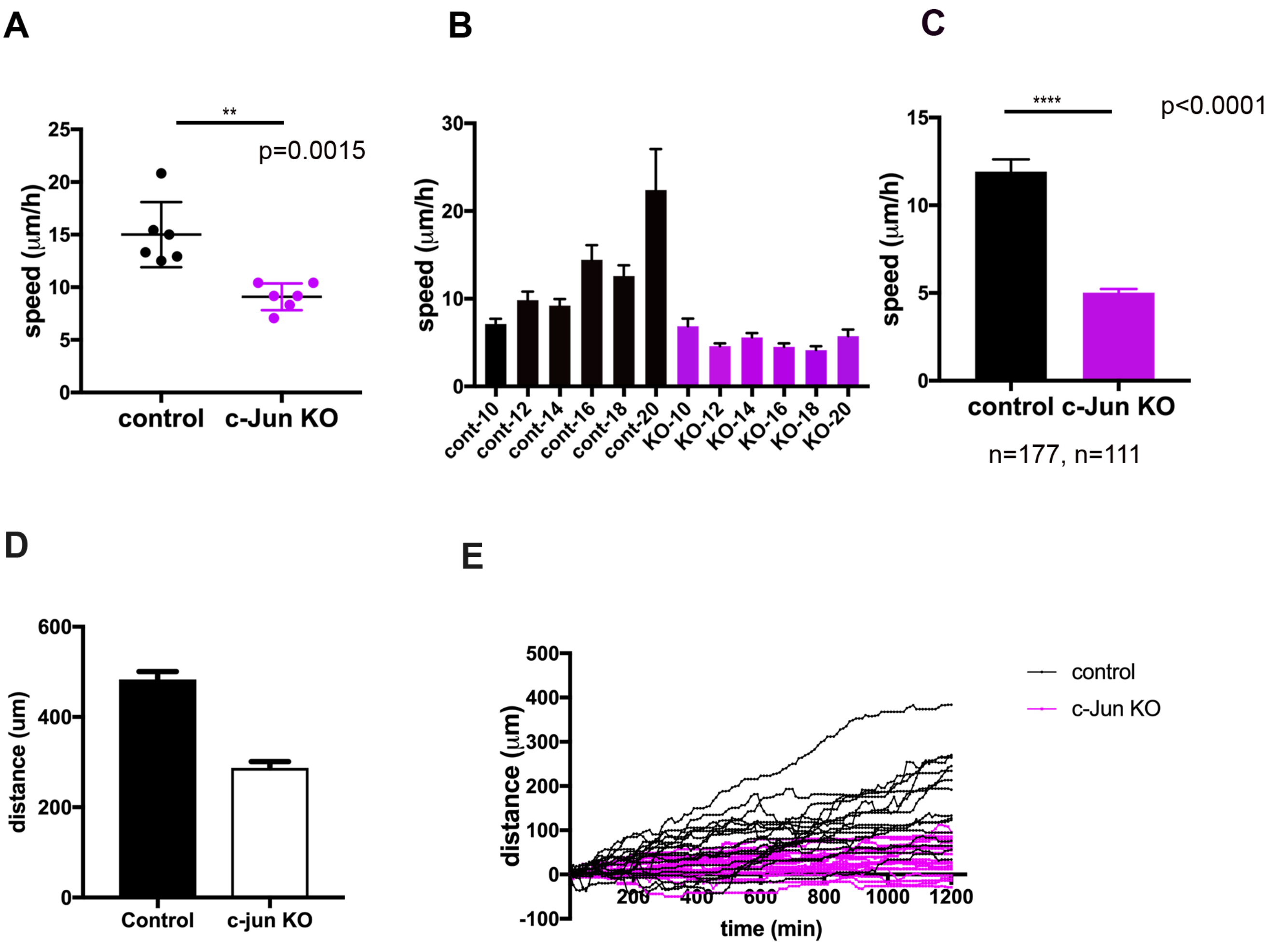


**Supplementary Fig. 5 | SC c-Jun supports both SC and cancer cell migration. A**, Quantification of control and c-Jun KO HEI-286-SC speed in two dimensions. **B**, Quantification of control and c-Jun KO HEI-286-SC speed in three dimensions in microchannels of sizes from 10 to 20 µm. **C**, Quantification of control and c-Jun KO HEI-286-SC speed in three-dimension. **D**, Quantification of length of columns formed by MiaPaCa-2 combined with WT HEI-286 SCs or with c-Jun KO HEI-286-SCs. **E**, Individual tracks of cancer cells in microchannels occupied by control vs. c-Jun KO HEI-286 SCs. (n=21 in each group).
