## Supplementary material for "Reprogrammed Schwann Cells Organize into Dynamic Tracks that Promote Pancreatic Cancer Invasion": sup Fig6

Sup Fig 6

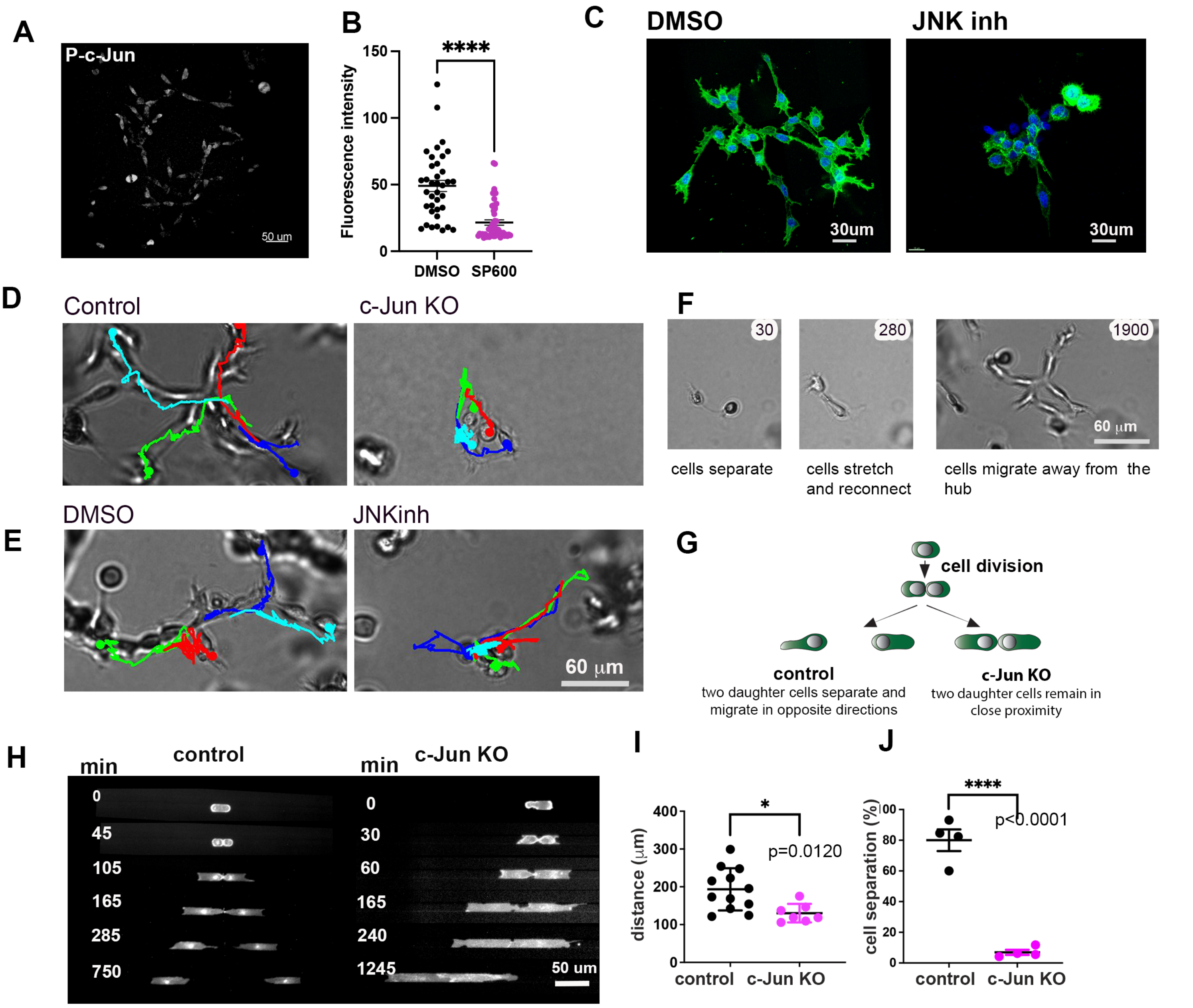


**Supplementary Fig. 6 | WT and c-Jun KO SCs behavior. A**, Confocal images of phospho-c-Jun (P-c-Jun) staining in HEI-286 GFP SCs in Matrigel. **B**, Effect of SP600125 on P-c-Jun expression in HEI-286 GFP SCs. **C**, Confocal images of HEI-GFP-SCs treated with JNKi SP600125 or DMSO in Matrigel showing lack of SC organization in JNKi-treated cells. **D-E**, Images of cells from 72h time-lapse movies of control and c-Jun KO HEI-286 SCs (D) and DMSO or JNK inhibitor treated HEI 286 SCs (E) grown in 3D Matrigel with colored cell tracking. **F**, Images of WT SCs from time-lapse movies showing formation of a branched organized structure in 3D: cells separate after division (1); cells stretch and reconnect, reestablishing contacts (2); cells divide, organize in a structure that spreads out (3). Time in minutes **G**, Schematic show the separation and positioning of two daughter HEI-286 SCs after mitosis in Matrigel is c-Jun-dependent. **H**, Time-lapse images of a dividing control and c-Jun-KO HEI-286 SCs and the two daughter cells in microchannels. **I**, Quantification of (H), the distance between two daughter cells following mitosis for control vs. c-Jun KO HEI-286 SCs in a 400 min time period (control n=16, c-Jun KO n=12, mean ± SEM). **J**, Quantification of percentage of daughter cells that separate following mitosis for control versus c-Jun KO HEI-286 SCs. (4 independent experiments, n>20 cells per condition in each experiment, mean ± SEM)
